# Glutamatergic Control Of A Pattern-Generating Central Nucleus In A Gymnotiform Fish

**DOI:** 10.1101/2020.10.05.326397

**Authors:** Virginia Comas, Michel Borde

**Author notes:** Corresponding author: Michel Borde.

## Abstract

The activity of pattern-generating networks (CPG) may change under the control exerted by various neurotransmitters and modulators to adapt its behavioral outputs to different environmental demands. Although the mechanisms underlying this control have been well established in invertebrates, most of their synaptic and cellular bases are not yet well understood in vertebrates. *Gymnotus omarorum*, a pulse-type gymnotiform electric fish, provides a well-suited vertebrate model to investigate these mechanisms. *G. omarorum* emits rhythmic and stereotyped electric organ discharges (EODs), which function in both perception and communication. The EOD is considered the behavioral output of an electromotor CPG which, modulated by descending influences, organizes adaptive electromotor behaviors in response to environmental and social challenges. The CPG is composed of electrotonically coupled intrinsic pacemaker cells, which pace the rhythm, and bulbospinal projecting relay cells that contribute to organize the pattern of the muscle-derived effector activation that produce the EOD. We used electrophysiological and pharmacological techniques in brainstem slices of *G. omarorum* to investigate the underpinnings of the fast transmitter control of its electromotor CPG. We demonstrate that pacemaker, but not relay cells, are endowed with ionotropic and metabotropic glutamate receptors subtypes. We also show, for the first time in gymnotiformes, that glutamatergic control of the CPG likely involves both AMPA-NMDA receptors transmitting *and* only-NMDA segregated synapses contacting pacemaker cells. Our data shed light on the fast neurotransmitter control of a vertebrate CPG that seems to exploit the kinetics of the involved postsynaptic receptors to command different behavioral outputs.

**NEW & NOTEWORTHY:** Underpinnings of neuromodulation of pattern-generating central networks (CPG) have been well characterized in many species. The effects of fast neurotransmitter systems remain, however, poorly understood. This research uses *in vitro* electrophysiological and pharmacological techniques to show that the neurotransmitter control of a vertebrate CPG in gymnotiform fish involve the convergence of only-NMDA and AMPA-NMDA glutamatergic synapses onto neurons that pace the rhythm. These inputs may organize different behavioral outputs according to their distinct functional properties.

## INTRODUCTION

The study of the neural strategies that organize behavior is a major challenge in Neuroscience. Advancing our knowledge in this field critically depends on the use of experimental paradigms that provide multiple levels of analysis (i.e., behavioral, network, cellular, synaptic, and molecular levels) as well as powerful techniques. Understanding the neural underpinning of behavior is further facilitated when there is a direct relationship between the neural mechanisms under study and the behaviors that result from these mechanisms. Pattern-generating neural systems (Central Pattern Generators, CPGs) are examples of valuable experimental models that meet this requirement (Katz 2016). Most CPGs can be considered as neural networks involving two interconnected functional levels (Dougherty and Ha 2019; McCrea and Rybak 2008). One level is represented by a central pacemaker that generates the rhythmic activity and the other level is composed by a set of neurons responsible for the specific patterned sequence of activation of peripheral effectors. In spite of the apparent stability of its neuronal components and intrinsic connectivity, extensive experimental evidence in invertebrates (Briggman and Kristan 2008; Marder et al. 2014; Puhl and Mesce 2008) and vertebrates (Bass 2008; Borde et al. 2020; Grillner and El Manira 2015, 2020; Steuer and Guertin 2019; Zornik and Kelley 2011) indicate that the activity of CPGs may present changes controlled by various neurotransmitters and modulators that modify the behavioral outputs to adjust to different environmental and social demands. Although the mechanisms underlying these modulations have been particularly well established in invertebrates, most of their synaptic and cellular basis in vertebrates are not yet well understood.

The pacemaker nucleus (PN) in gymnotiform electric fish is considered a two-layered model CPG (Borde et al. 2020) that provide an advantageous model system to investigate the direct impact of CPG modulations on an overt behavior. The rhythmic layer of this CPG is represented by a group of pacemaker cells (PM-cells) and the patterning layer comprises the projecting relay cells (R-cells) and their synaptic targets at the spinal cord; i.e., a topographically distributed population of electromotor neurons that innervate the effector. Under the control of the PN, these species emit rhythmic and stereotyped electric organ discharges (EOD) that allows electric fish to detect physical properties of their nearby environment and to communicate among conspecifics (Caputi et al. 2005; Caputi and Budelli 2006; Lewis 2014; Lissmann 1951; Lissmann and Machin 1958; Lorenzo et al. 2006). PM- and R-cells aggregate at the midline near the ventral surface of the medulla and form the PN (Fig. 1A). In pulse-type gymnotiforms, PM-cells are grouped dorsally and set the pace of the EOD whereas R-cells occupy the ventral aspects of the PN (Fig. 3A). PM-cells are electrotonically coupled with each other and with R-cells (Bennett 1971; Bennett et al. 1967) and they form a rather simple circuit with exclusive feedforward connections. The command for each EOD is thus initiated at PM-cells that discharge synchronously and then transmitted 1:1 to R-cells that ultimately translate the command drive to the electric organ, producing a species-specific EOD waveform pattern (Curti et al. 2006; Lissman 1951; reviewed in Caputi et al. 2005).

**Figure 1.**
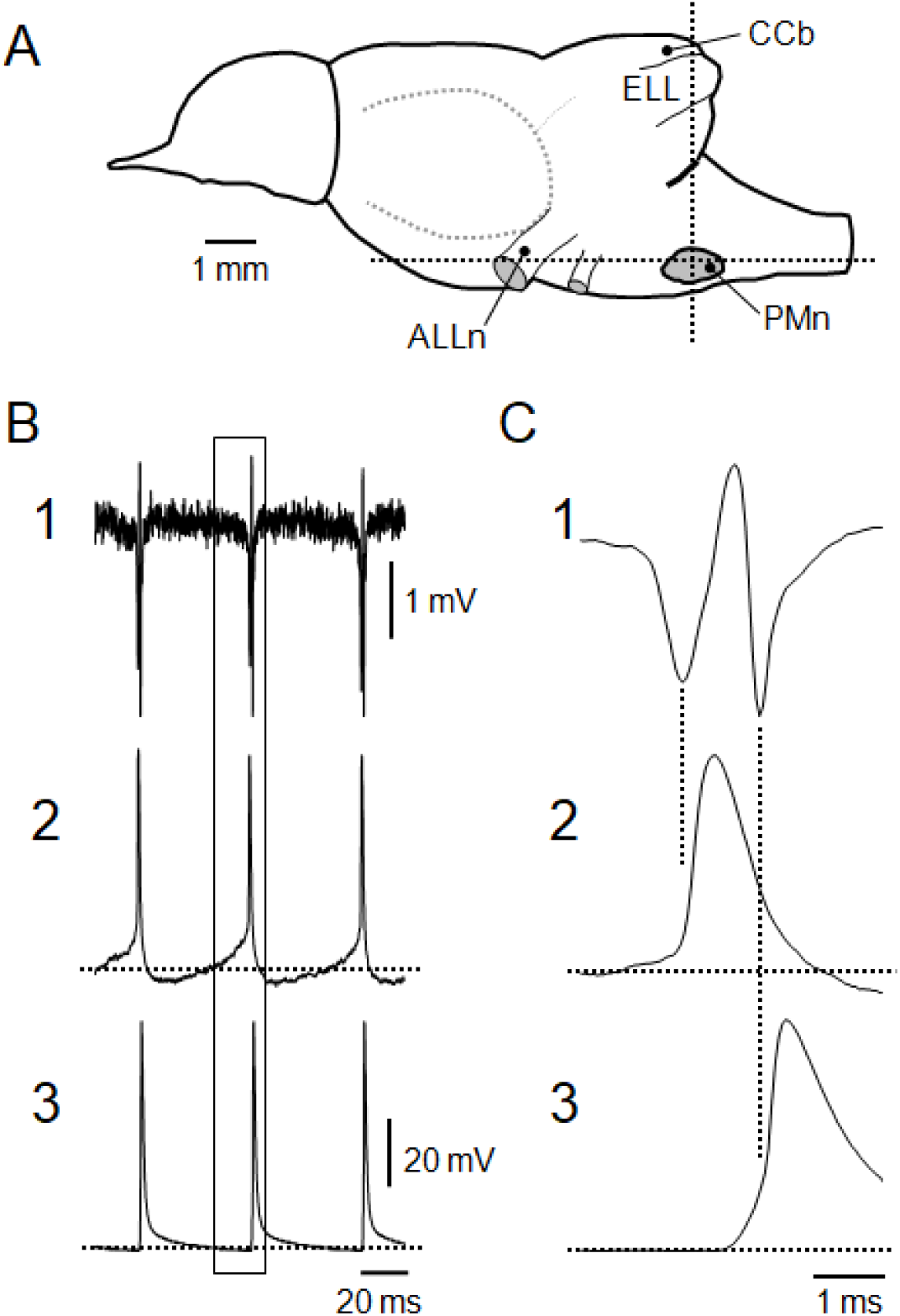
Rhythmic activity of the PN *in vitro*. **A.** Schematic drawing of a lateral view of the brain of *Gymnotus omarorum*. Anterior lateral line nerve (ALLn), corpus cerebelli (CCb), electrosensory lateral line lobe (ELL) and the approximate location of the PN were included in the scheme as references. The dotted lines represent the levels at which the brainstem was sectioned to obtain 400 µm thick transverse (vertical line) and horizontal (horizontal line) slices containing the PN. **B**. Recordings of the PN spontaneous rhythmic activity in a transverse slice. **B1.** Field potential recording of the PN obtained near PM-cells. **B2.** Intracellular recording of a PM-cell. **B3**. Intracellular recording of a R-cell. Paired recordings of the PN field potential and of the transmembrane potential of either a PM- or a R-cell were obtained and traces were aligned taking the field potential of each pair as a temporal reference. **C.** Details of discharges shown in B (boxed region) displayed at a faster sweep speed showing the temporal correlation between extracellular and intracellular events. **C1.** Field potential. **C2.** PM-cell spike. **C3.** R-cell spike. The peak of the first and second negativity of the PN spontaneous field potential correlates with the rising phase of the action potential of the PM- and the R-cell (vertical dotted lines). In B and C, a membrane potential of -70 mV is indicated in traces of 2 and 3 by dotted lines.

EOD modulations observed during specific adaptive and social behaviors constitute accessible, prominent, and distinctive electromotor behaviors. These depend on the activation of specific descending inputs to the PN, that arise from distinct prepacemaker nuclei (PPn; Caputi et al. 2005; Comas and Borde 2010; Comas et al. 2019). More recently, hormonal and peptidergic influences upon the electromotor system have also been characterized (reviewed in Silva 2019). Electromotor behaviors include a wide variety of changes in the rate or in the waveform of the EOD. A wealth of previous studies (Caputi et al. 2005; Kawasaki and Heiligenberg 1989, 1990; Lorenzo et al. 2006; Quintana et al. 2011, 2014; Spiro 1997) show that distinct types of short term rate EOD modulations depend not only on the specific PPn being activated, but also on both the cellular target of PPn axons within the PN and the neurotransmitter receptors involved (Caputi et al. 2005; Curti et al. 1999; Dye et al. 1989; Kawasaki and Heiligenberg 1989, 1990; Keller et al. 1991; Quintana et al. 2011, 2014). More recently, the modulation of electrophysiological properties of PN neurons has emerged as a novel putative mechanism, endowing electromotor networks with higher functional versatility (Comas et al. 2019). Activation of GABAergic inputs to PM-cells induce them to silence their discharge provoking EOD interruptions (Comas et al. 2019; Kawasaki and Heiligenberg 1989, 1990). Glutamatergic inputs to these cells, in turn, evoke slow and long lasting rises in EOD rate *via* the activation of NMDA receptors (NMDAR) by NMDAR-only synapses (Curti et al. 1999, 2006; Juranek and Metzner 1997, 1998; Kawasaki and Heiligenberg 1988, 1989, 1990; Kawasaki et al. 1988; Keller et al. 1991; Spiro et al. 1994). In pulse type gymnotiformes an increase in EOD rate in responses to AMPA receptor (AMPAR) agonists applied to PM-cells has been described (Curti et al. 1999; Quintana et al. 2014). Nevertheless, electromotor behaviors derived from activation of those receptors as well as the pattern of activation by glutamatergic descending inputs are still unknown. In R-cells, GABAergic innervation is apparently lacking (Comas et al. 2019; Kawasaki and Heiligenberg 1990; Kennedy and Heiligenberg 1994). However, as described in several gymnotiform species, these cells are contacted by glutamatergic inputs whose activation triggers suprathreshold R-cells depolarizations, emitting the bursts of high rate small discharges called chirps (via AMPAR) or abruptly stopping the EOD (via NMDAR) (Caputi et al. 2005; Kawasaki and Heiligenberg 1989, 1990; Quintana et al. 2011, 2014; Spiro 1997).

Although the role of fast neurotransmitter modulations of the electromotor CPG exert on electric behavior is indisputable, several important questions remain unsolved. For example, can AMPAR of PM-cells be activated by the synaptically released glutamate? Do they participate in AMPAR-only synapses or they are involved in classical AMPAR-NMDAR transmitting synaptic contacts (Clark and Cull-Candy 2002; Cottrell et al. 2000; Traynelis et al. 2010)? As only-NMDA synapses have been proposed to involve receptors with slow kinetics likely containing subunits of the GluN2B type (Nakayama et al. 2005; Paoletti 2011), are similar receptors involved in the control of the electromotor CPG? To address these issues, in the present study we developed an *in vitro* preparation of *Gymnotus omarorum* brainstem slices containing the PN. Among gymnotiform fish, this species is more suitable for our study since the rhythmic layer of the electromotor CPG is apparently the exclusive cellular target of the descending glutamatergic inputs (Curti et al. 1999, 2006). This preparation allows the extra- and intracellular recording of responses of PM- and R-cells to: i, glutamate and its agonists for different receptor subtypes applied juxtacellularly and ii, glutamate released by electric activation of descending inputs to the PN. We show that PM-cells, but not R-cells, are endowed with functional NMDA, AMPA and metabotropic glutamate receptors and that NMDAR containing subunits of the GluN2B type are not involved in the control of the electromotor CPG. We also demonstrate for the first time in gymnotiformes that glutamate released at certain synaptic terminals co-activates AMPAR and NMDAR that are most probably co-localized at glutamatergic synapses onto PM-cells. Our study contributes to a better understanding of how synaptic inputs to a vertebrate pattern-generating central nucleus may generate adaptive behavioral outputs.

## MATERIALS AND METHODS

### Preparation of slices

*Gymnotus omarorum* (n=32, 11-16 cm in length) were collected from Laguna del Sauce (34°51’S, 55°07’W, Department of Maldonado, Uruguay). All the experimental procedures were conducted in accordance to the guidelines set forth by the Comisión Nacional de Experimentación Animal (exp. 071140-000092-13).

Individuals were anesthetized and placed in a plastic chamber on a wet sponge. All surgical areas and fixation points were heavily infiltrated with 2% Lidocaine hydrochloride. During surgical procedures, the head was maintained in a horizontal position by a pair of plastic tipped metal bars attached to the box and the gills were perfused with MS222 dissolved in iced tap water (0.3 mg/l). The dorsal surface of the brain was exposed while bathed with cold Na-free artificial cerebrospinal fluid (ACSF)- sucrose solution (see below). The brain with part of the spinal cord was rapidly removed from the skull and submerged in cold ACSF-sucrose solution. Transverse and horizontal sections (Fig. 1A) of the brainstem (400 µm thick) containing the PN were obtained under cold ACSF-sucrose using a Vibratome 1000 plus (The Vibratome Company, Saint Louis, MO, USA), and were incubated (>1h, at room temperature, 21-23 °C) in a 1:1 solution of ASCF-sucrose and control ACSF solution (see below). Slices were transferred to a 2 ml recording chamber fixed to an upright microscope stage NIKON (Eclipse FN1, Nikon Company, Minato, Tokio, Japan) equipped with infrared differential interference contrast (DIC) video microscopy and a 40x water immersion objective. Slices were perfused with carbogen-bubbled ACSF (1.5-3 ml/min) and maintained at room temperature (20-23 °C). Under these experimental conditions, the PN maintains its spontaneous synchronized activity with a stable firing rate for at least 6 hours (Fig. 1B).

### Recording and stimulation

The electric activity of the PN was monitored using an Axoclamp 2B amplifier (Molecular Devices, San José, CA, USA) by means of extracellular recordings of field potentials or by recording intracellularly from PM- or R-cells. For extracellular recordings, patch pipettes (4-8 MΩ) filled with ACSF were usually placed near R-cell somas. Intracellular recordings were performed with a patch pipette in the whole cell configuration from the soma of the cells. The patch pipette (5-10 MΩ) was filled with a potassium gluconate-based intracellular solution (see below). In some experiments (n=14) for intracellular recordings, sharp electrodes were used (4M potassium acetate, 60-140 MΩ). Microelectrodes were placed under visual control with a hydraulic micromanipulator (Narishige, Setagaya, Tokyo, Japan). Signals were low-pass filtered at 3.0 kHz, sampled at 10.0 kHz through a Digidata 1322A (Molecular Devices, San José, CA, USA) and stored in a PC computer for further analysis with the aid of the pClamp programs (Molecular Devices, San José, California, USA).

Intracellular recordings of electrical activity of PN neurons were performed under current clamp. Long lasting depolarizing current pulses (800-1000 ms, 2-4 nA) were injected to explore repetitive firing behavior in both cellular types.

In horizontal brainstem slices, glutamatergic PN afferent fibers were electrically stimulated. Guided by results obtained in previous studies in *G. omarorum* (Comas and Borde 2010), a bipolar stimulating electrode (Nichrome, 75 µm) was placed near the midline over the medial longitudinal fasciculus at a distance of 200 - 400 µm from the PN in the rostral direction. The ventral portion of this fasciculus is thought to contain descending axons of PPn neurons. Stimuli consisted in short trains of 3 pulses (3 - 6 V; 0.02 ms each) at 250 Hz mimicking the discharge of putative PPn neurons responsible for the acceleration of the EOD triggered by the activation of the command neuron for escape behavior (Comas 2010). Trains were repeated every 10s. Strength of stimulation was adjusted to produce transient accelerations of the PN discharge rate which were very similar in amplitude and time course to these EOD accelerations (Curti et al. 1999, 2006). Consistently the time elapsed between the spike of either PM or R-cells immediately preceding the stimulus and the moment of stimulation was kept at approximately 30% of the duration of the control interval. The interval during which the stimulus was applied was called I_0_, and the subsequent interval was designated I_1_. The interval that immediately preceded the stimulation was designated as the “control” interval (or I_C_) (see Fig. 6A).

Micropipettes filled with different solutions of glutamate or its agonists were used to pressure ejection of drugs in transversal slices at the PN (Picospritzer II; General Valve, NY, USA). Pulses of 20 psi and 20-150 ms duration ejected relatively small volumes (5–15 pl) that were calculated from the diameter of microdroplets estimated by inspection of the area of mechanical distortion of tissue provoked by the ejected solution under the 40x magnification. Ejection of microvolumes was repeated 5 to 10 times with a 20 s interval. The use of small volumes was preferred in order to avoid affecting responses with mechanical artifacts (however, see below). Volumes of drugs were selected in order to evoke mild accelerations (range 1 - 12.5 Hz). The tips of these micropipettes were positioned at different locations within the PN closely to PM- or R-cells somas under visual control (at distances ranging from 30 to 70 µm). In most cases, within the same experiment, similar microvolumes of drug-containing solutions were applied juxtacellularly to PM- and R-cells. In selected experiments, control of drug effects were made by ejecting similar volumes of ACSF at both PM and R-cells. In some rate responses to drugs applied to PM-cells, small and brief sharp increases in the rate coinciding with the pressure delivery of drug-containing solutions were observed. These responses, which were interpreted as being produced by mechanical stimulation of the cells, were still observed in the presence of glutamatergic antagonists and could also be provoked by the injection of ACSF (not shown).

### Analysis of responses

The Clampfit software (pClamp 10) was used to analyze the effect of stimuli on the rate of the spontaneous discharge of the PN. Individual discharges (field potential or intracellular recorded spike) were detected (threshold search) and instantaneous frequency versus time plots of events were constructed. Rate responses were averaged, and peak amplitude was calculated by substracting the basal spontaneous rate to the highest frequency attained during the responses. The time course of rate responses was quantified by measuring the following parameters: the rise and decay time constants (τ_RISE_ and τ_DEC,_ respectively) by adjusting a single exponential in the rising and the decay phases of responses, and their duration at the half amplitude (half width, HW).

The absence of evident effects of agonists applied to R-cells was confirmed by comparing the mean PN discharge rate before and after the ejection. For this purpose the rate was averaged for a period of 1 s preceding the stimulus and a period of 5.5 s after the ejection. The duration of the post-ejection period for this analysis was taken based on the typical time course of the rate responses to similar microvolumes of glutamate applied to PM-cells (mean duration of 5.5 s, see Results section).

In intracellular recording of R-cells, subthreshold membrane potential changes evoked by agonist ejections were explored by comparing the mean minimum potential level (MPL; Fig. 3B) measured before (1s duration period) and after (5.5s duration period) the ejection.

### Estimation of NMDA and AMPA components of responses

The relative contribution of the NMDAR and AMPAR to responses to glutamate ejection or afferent stimulation was estimated by measuring components of the response due to activation of each receptor subtype (NMDA_comp_ and AMPA_comp_). For responses evoked by PPn axon stimulation, the magnitude of each component was estimated by a point-by-point subtraction of responses obtained before and during the perfusion of a selective glutamate receptor antagonist (CNQX or AP5, see for an example Fig. 6). For agonist-induced rate responses, peak amplitude of each component was calculated by subtracting the peak amplitude of the responses evoked before and during perfusion of the specific blocker. In both cases, for estimation of the NMDA receptor-dependent component the formula was:

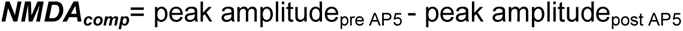

and for the AMPA receptor-dependent component,

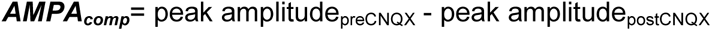

Once the magnitude of each component was obtained, the NMDA to AMPA ratio of responses was calculated as:

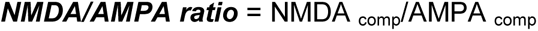

Usually, CNQX and AP5 were added sequentially to the perfusion solution (as illustrated in Fig. 7 for example) in such a way that rate responses were evaluated under one or both glutamatergic blockers. The order in which they were added to the perfusion solution (with either CNQX or AP5 as the first blocker applied) depended on the specific experimental design.

The dependence of the NMDA/AMPA ratio on the order of administration of blockers was estimated by calculating the order dependence index (OD index) as follows:

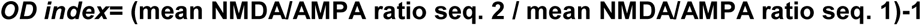

where seq. 1 corresponds to the CNQX→CNQX+AP5 sequence when AP5 was added to the ACSF containing CNQX and seq. 2 to the AP5→AP5+CNQX sequence obtained by adding CNQX to a solution that contains AP5. If the mean NMDA/AMPA ratio is independent of the sequence:

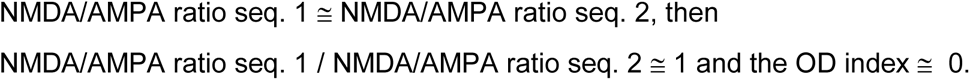

### Drugs and solutions

ACSF solution contained (in mM): 124 NaCl, 3 KCl, 0.75 KH_2_PO_4_, 1.2 MgSO_4_, 24 NaHCO_3_, 10 D-Glucose, 1.6 CaCl_2_, pH 7.2-7.4 after saturation with carbogen (Spiro 1997), while NaCl was substituted with 213 sucrose in ACSF-sucrose solution. Mg^2+^ free solution was obtained by substituting the MgSO_4_ (1.2 mM) of the ACSF-solution with CaCl_2_, increasing its concentration to 2.8 mM. Intracellular solution in patch pipettes contained (in mM): 140 K-gluconate, 0.2 EGTA, 4 ATP-Mg, 10 HEPES, pH 7.3). Effects of the following substances applied juxtacellularly (dissolved in ACSF solution) were investigated: L-glutamic acid (glutamate, 10 mM), NMDA (50 μM), AMPA (50 μM), and trans-(±)-1-amino-1,3-cyclopentanedicarboxylic acid (trans-ACPD, 500 μM), an agonist for metabotropic glutamate receptors (mGluR). Glutamatergic antagonists (±)-2-amino-5-phosphonopentanoic acid (AP5; 50 μM), ifenprodil (10 μm), 6-cyano-7-nitroquinoxaline-2,3-dione (CNQX, 20 μM), and (±)-amino-4-carboxy-methyl-phenylacetic acid (MCPG, 600 μM) for NMDAR, NMDAR containing GluN2B subunits, AMPAR and mGluR subtypes, respectively, also were dissolved in the ACSF solution, and perfused during at least 30 minutes. During afferent stimulation experiments, ACSF contained picrotoxin (100 µM) to block GABAergic inputs to the PN that could be recruited by the electric stimuli. These substances were obtained from Sigma-Aldrich (Saint Louis, MO, USA) or Tocris Bioscience (Bristol, UK).

### Statistical analysis

All data are presented as mean ± standard deviation (SD) and were statistically processed using the Past software (Hammer et al. 2001). Once subjected to the Shapiro-Wilk normality test, Student *t-*test and paired *t*-test were used for independent and paired variables, respectively. For data sets with non-normal distribution, the non-parametric statistical tests Mann–Whitney *U* test and Wilcoxon test, for independent and paired variables, respectively, were used. To preserve visibility of the figure panels, significance levels of <0.05 and <0.01 are marked by one or two asterisks, respectively. The absence of statistical significance is indicated by n.s. (non significant). Along the text, absolute *P* values are reported. For estimation of SD of the OD index by computing the mean±SD of NMDA/AMPA ratios as input values, the Gaussian algorithm of error propagation of a quotient was used.

## RESULTS

The PN was spontaneously active *in vitro* and discharged rhythmically with a mean rate of 18.5±3.7 Hz, values which falls within the range of EOD rates observed in resting behaving diurnal fish (15-25 Hz). A representative example of simultaneous recordings of the spontaneous extracellular field potential and intracellular activity of either a PM- or a R-cell is shown in figure 1B. The temporal correlation of the recorded electric events (Fig. 1B, C) shows that the normal sequence of activation is conserved *in vitro*, i.e.: synchronous activation of PM-cell drives the synchronous discharge of R-cell in a 1 to 1 manner. The amplitude and waveform of field potentials recorded near PM-cells (Fig. 1C1) and the timing of PM- and R-cell activity is also preserved when compared to *in vivo* results. In addition, the spike of PM-cells preceded the spike of R-cells in approximately 0.9 ms, a similar delay to that reported *in vivo* (Curti et al 2006). In intracellular recordings, PM-cells showed an unstable membrane potential with a prolonged “pacemaker potential”, a slow depolarization preceding the spike onset (Fig. 1B2). The action potential of R-cells, in contrast, rose abruptly from the baseline membrane potential (Fig. 1B3). Characteristically both types of neurons also differed in their responses to long lasting depolarizing suprathreshold pulses (Fig. 2). PM-cells usually responded with a single action potential at the onset of the current pulse (arrowhead in Fig. 2A) irrespective of its amplitude. In contrast, suprathreshold depolarizations of R-cells evoked repetitive high rate small amplitude discharges without any detectable frequency adaptation (Fig. 2B). The above data indicate that the *in vitro* slice preparation preserves the electrophysiological properties of PM- and R-cells and intrinsic network organization of the PN.

**Figure 2.**
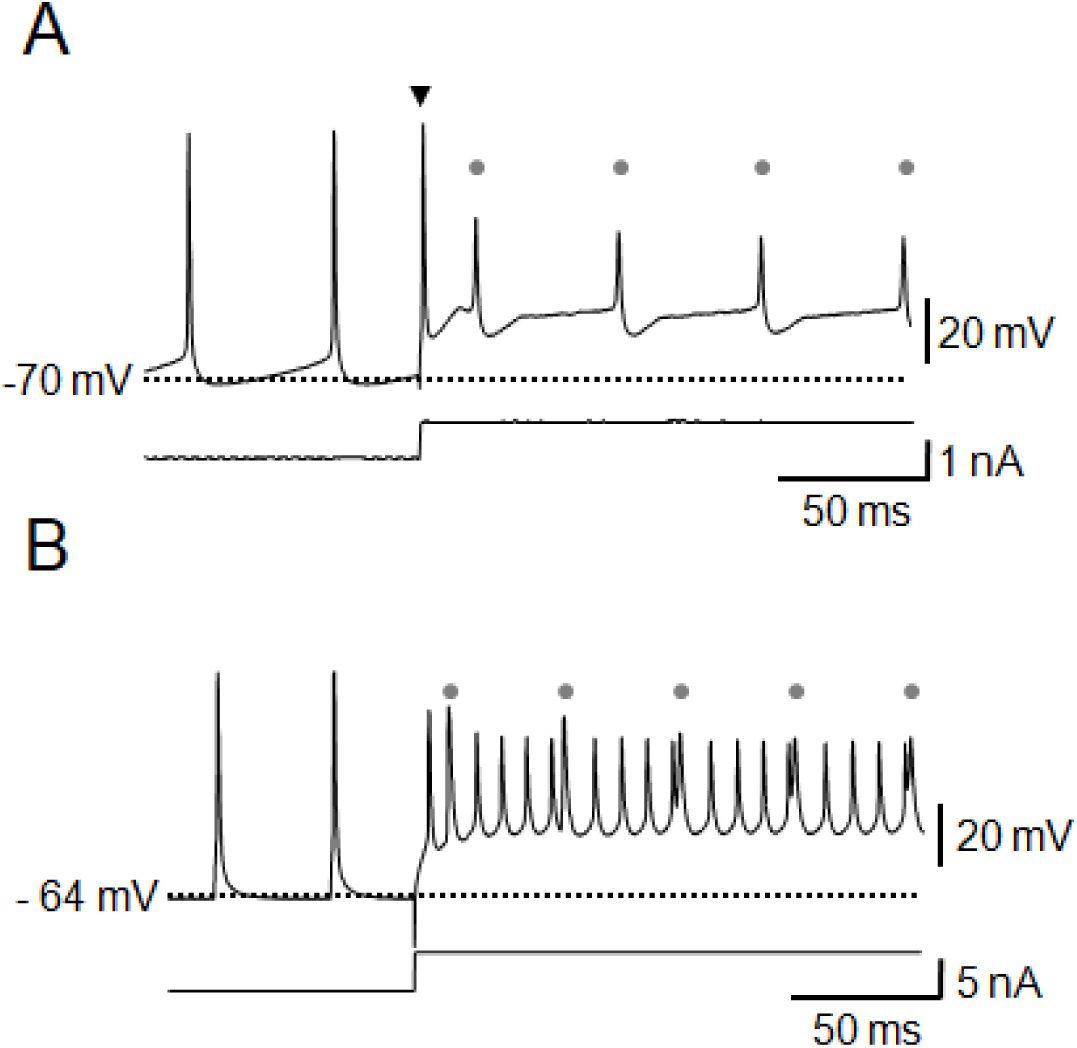
Main electrophysiological features of PN neurons. **A.** Intracellular recording of a PM-cell obtained before and during a long lasting depolarizing current step (bottom trace). Before current injection, spontaneous rhythmic action potentials are preceded by a slow depolarizing trend (pacemaker potential). Typically, suprathreshold current pulses evoke a single action potential at the onset of the stimulus (black arrowhead) while, although reduced in amplitude and with altered waveform, the regular rhythmic action potential can be still observed during the depolarization (gray dots) discharging at almost the same pre-ejection rate. **B.** Same as A but recording was obtained from a R-cell. Spontaneous pre-ejection action potentials arise abruptly from a stable basal membrane potential. Depolarization evokes the repetitive discharge of low amplitude action potentials at a relatively high rate superimposed to the ongoing regular rhythmic discharge driven by pacemaker neurons (gray dots). Numbers at the left of each recording show the membrane potential of cells indicated by the dotted lines.

### Effects of glutamate agonists on R-cells

In this series of experiments, the sensitivity of R-cells to juxtacellular application of glutamate, AMPA and NMDA (Fig. 3A) was assessed. In addition to spike rate, the presence of subthreshold responses to ionotropic glutamate receptor agonists was specifically explored by recording the membrane potential of R-cells during agonist application.

**Figure 3.**
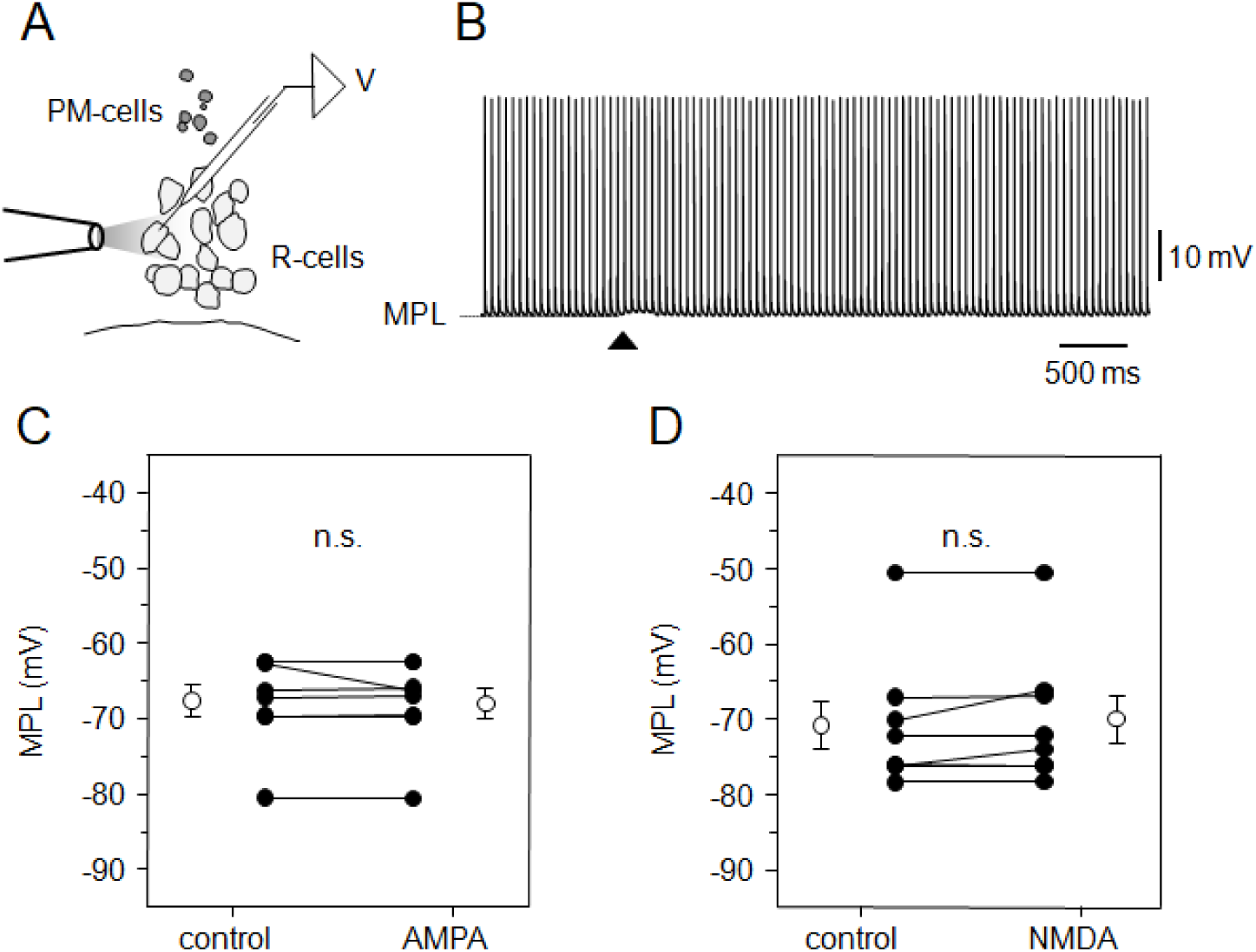
Effects of pressure-applied glutamate agonists on R-cells. **A.** Schematic drawing of a transverse section of the PN illustrating a drug-containing micropipette for pressure ejections (at left), the somas of PM- and R-cells and the location of the recording electrode (V). **B.** Intracellular recording of a R-cell before and during the response to glutamate ejection (10 mM, 20 ms, 20 psi, black arrowhead). MPL at left indicates the minimum potential level before Glu ejection. **C.** Connected dot plots of the averaged MPL before (control) and after the juxtacellular ejection of a microvolume of AMPA (50 µM). Each dot pair represents a particular experiment. Mean ± SD of MPL values are represented by white circles (n=8; Wilcoxon test; p=0.547)**. D.** Same as C but illustrating the effect of NMDA (50 µM, n=8; Wilcoxon test; p=0.148).

Figure 3B illustrates a representative recording of an R-cell obtained before and after the application of a microvolume of glutamate. In the example, there were no evident changes either in the MPL or in the spike rate after glutamate application. However in some experiments (4 out of 6 experiments), a transient and small (at most 0.8 mV) depolarizing change in the MPL was observed. The mean MPL measured before and after glutamate applications was -73.2±2.9 mV and -72.9±3.1 mV, respectively (n=6, paired *t* test, p=0.027). Although these values were slightly different, in 2 experiments the small change in the MPL provoked by glutamate persisted after 30 minutes of bath application of glutamate receptor blockers suggesting its artifactual nature (not shown). The spike rate of R-cells was not modified by glutamate being 20.2±1.8 Hz and 20.2±1.8 Hz, before and after injection, respectively (n=6, paired *t* test, p=0.055). In line with these results, neither AMPA nor NMDA injections provoked changes in the MPL (Fig. 3C, D). The mean MPL measured before and after agonists applications were -67.6±2.1 mV and -68.0±2.0 mV, respectively (n=8, Wilcoxon test, p=0.547) for AMPA and -70.9±3.2 mV and -70.0±3.2 mV, respectively (n=8, Wilcoxon test, p=0.148) for NMDA. Spike rate was not affected by agonists being 29.4±1.4 Hz and 29.4±1.4 Hz before and after ejections of AMPA, respectively (n=8, Wilcoxon test, p=0.461) and 20.7±1.5 Hz and 21.0±1.6 Hz, before and after ejections of NMDA, respectively (n=8, Wilcoxon test, p=0.063).

### Effects of glutamate agonists on PM-cells

Juxtacellular application of microvolumes of glutamate and its agonists to PM-cells systematically provoked transient accelerations (mean duration 5.5±3.4 s) of PN discharges (Fig. 4A). Glutamate-evoked accelerations were in average 8.79±3.2 Hz (n=10) which correspond to an increase of 35.21±5.67 % relative to the basal pre-ejection rates and consistently associated with slow, low amplitude depolarization of PM-cell’s MPL of at most 2.3 mV (1.34±0.77 mV) (Fig. 4B, inset). Representative examples of PN rate responses evoked by pulse application of ionotropic and metabotropic glutamate receptors specific agonists to PM-cells are illustrated in Fig. 4C-E. AMPA evoked rate increases of 2.1±1.1 Hz (n=8) which represents an increase of 10.9±4.4 % relative to the basal pre-ejection rate (Fig. 4C). AMPA rate responses were reduced by 98.7 % ±2.47 (n=8) by CNQX (20 μM, Fig. 4C, gray circles). Rate responses to NMDA (Fig. 4D) reached peak amplitudes of 3.0±1.9 Hz (n=10) which correspond to an increase of 14.7±10.3 % of the basal pre-ejection rate. Bath application of AP5 (50 µM) reduced the rate responses to NMDA by 47.5±29.7 % (n=7, Fig. 4D, gray circles). The rate responses to NMDA were characteristically slower than those evoked by AMPA. Rise time constants of the responses to NMDA were significantly higher than to AMPA (NMDA; 356.8±78.0 ms, n=8; AMPA 199.6±100.9 ms, n=7; Wilcoxon test, p=0.015). Decay time constants of the NMDA responses were also significantly greater than those of the AMPA responses (NMDA; 7511.09±3954.19 ms, n=8; AMPA 3705.60±2626.21 ms, n=7; Student *t* test, p=0.04). However, although responses to NMDA were consistently slower than to AMPA their half width were not statistically different (NMDA; 6648.2±5577.93 ms, n=8; AMPA 3125.4±2077.88 ms, n=7; Mann Whitney *U* test, p=0.195)

**Figure 4.**
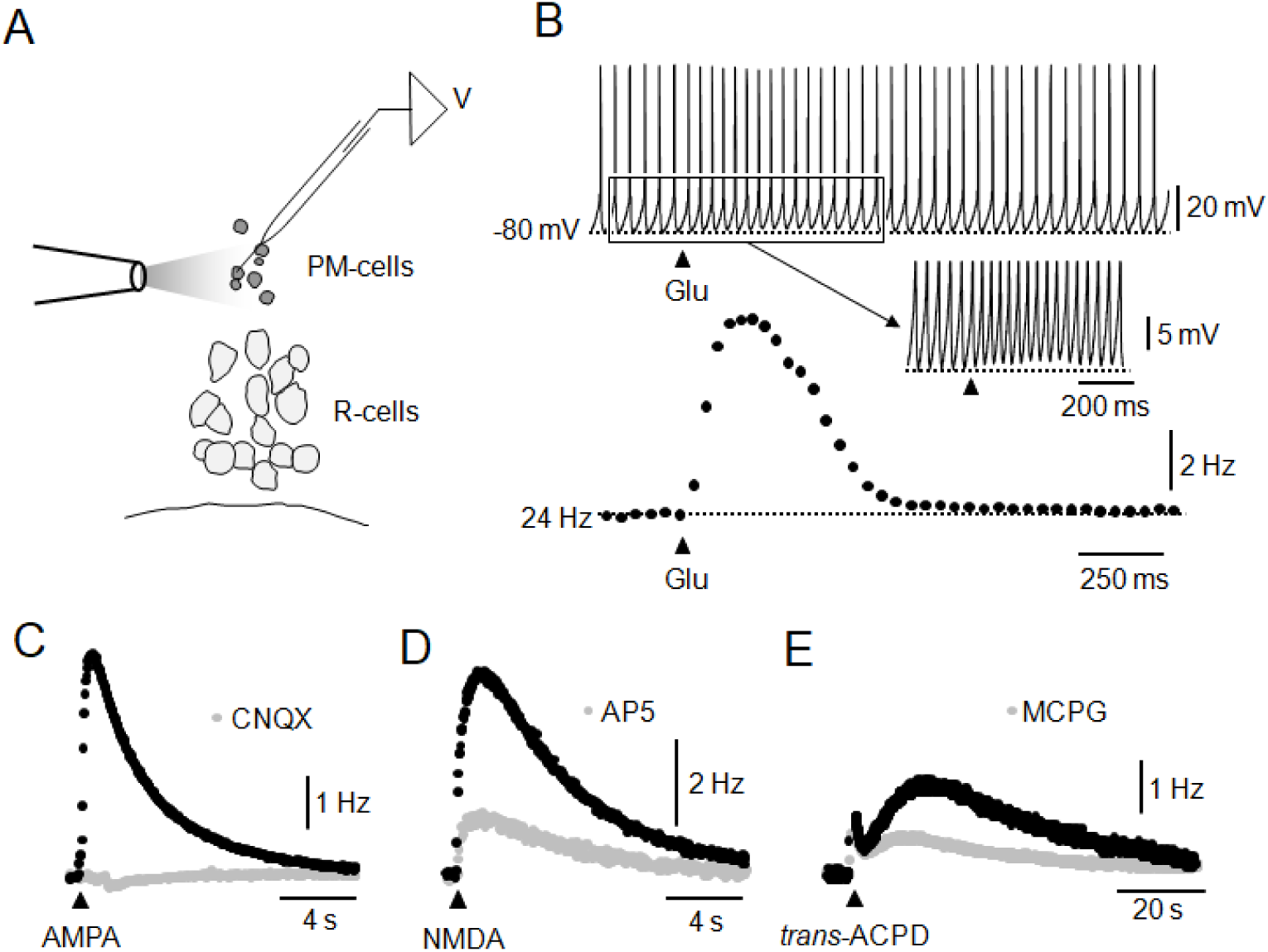
Effects of pressure-applied glutamate agonists on PM-cells. **A.** Schematic drawing of a transverse section of the PN illustrating a drug-containing micropipette for pressure ejections (at left), the somas of PM- and R-cells and a micropipette for intracellular recording of a PM-cell (V). **B.** Changes in the membrane potential and in the discharge rate of PM-cells in response to glutamate. **Top.** Intracellular recording of a PM-cell before and after the juxtacellular ejection of a microvolume of glutamate (10 mM, 20 psi, 20 ms, black arrowhead) obtained according to A. The number at left indicates the membrane potential indicated by the dotted line. Part of the response (boxed region) is illustrated in the inset below at a higher gain and a slower sweep speed to better illustrate the small depolarization that accompanies the acceleration of discharges. **Bottom**. Instantaneous frequency vs time plot of PM-cell discharges recorded in the top trace. Number at the left of the plot indicates the basal pre-ejection PN rate indicated by the dotted line. **C.** Rate responses of PN discharges before and after juxtacellular application of a microvolume of AMPA (50 µM, 150 ms, 20 psi) in ASCF (black circles) and after the perfusion of ACSF containing CNQX (20 μM, gray circles). **D.** Same as C but applying NMDA (50 μM, 200 ms, 20 psi). The response in ACSF (black circles) and after adding AP5 (50 μM, gray circles) to the ACSF are illustrated. **E.** Same as C but applying trans-ACPD (500 µM, 20 psi, 130 ms). The response in ACSF (black circles) and under perfusion of MCPG solution (600 μM, gray circles) are depicted. In C, D and E the black arrowhead indicates the moment of ejection of drugs. B, C, D and E are from different experiments.

Metabotropic glutamate receptors (mGluR) of PM-cells have been described *in vivo* as modulators of the PN discharge rate in *G. omarorum* and their contribution to the organization of slow and long lasting EOD modulations has been suggested (Curti et al., 1999). Juxtacellular ejection of microvolumes of the mGluR agonist *trans*-ACPD (500 μM) to PM-cells characteristically caused slow and long lasting increases in PN discharge rate (Fig. 4E) that closely resembled those observed *in vivo*. The rate response exhibited two components: a relatively fast and early peak followed by a delayed sustained component (∼1 Hz, 70 s; n=3). Under MCPG (600 mM), a broad spectrum mGluR antagonist, the slower component of the rate responses was partially blocked and their peak amplitudes were reduced by ∼53 % of control responses (n=2, Fig. 4E, gray circles).

To test whether only-NMDA synapses on PM-cells implicated in several well known electromotor behaviors involve receptors containing subunits of the GluN2B type, we explored the effect of ifenprodil (10 µM), a specific blocker of this type of NMDARs, on the rate responses to glutamate (Fig. 5). As illustrated in the example in Fig. 5A, average peak amplitudes and decay time constants were practically unaffected by ifenprodil, being 97.1±9.3 % and 102.9±16.1 %, respectively, from control rate responses (Fig. 5B). These values that were not different from control (Wilcoxon test, n=6, p=0.563 and p=0.25 for amplitude and decay time constant, respectively). In spite of this lack of effect, ifenprodil produced robust and reversible modifications of other variables: overall field potential amplitude was decreased, amplitude of action potential recorded intracellularly was reduced, and PN rate was slightly but consistently decreased (3.0±3.1 %). All changes were reverted after a 90-120 minutes washout.

**Figure 5.**
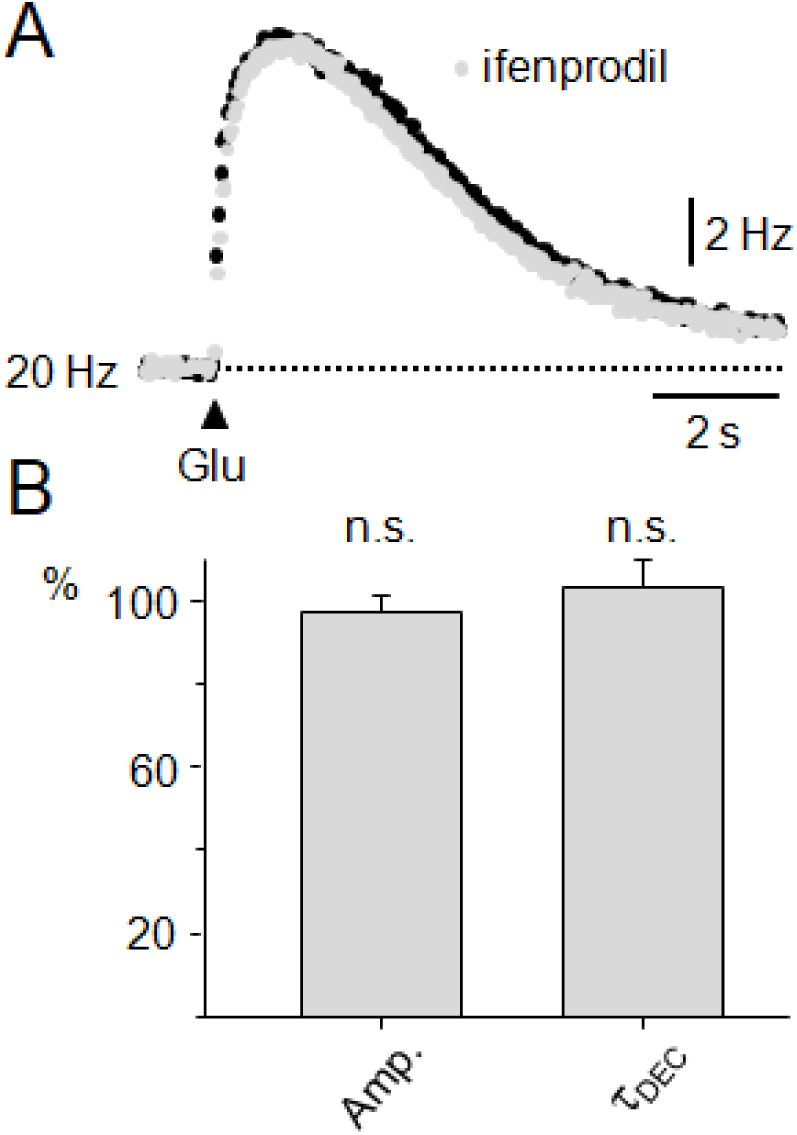
Effects of ifenprodil on rate responses to pressure-applied glutamate. **A.** Rate responses to glutamate (10 mM, 20 psi, 60ms, black arrowhead) obtained in ACSF (black circles) and after 15 minutes under ifenprodil (10 µM, gray circles). **B.** Summary data (bar chart, mean ± SD, n=6) of the effects of ifenprodil on the peak amplitude (Amp.) and the decay time constant (Ʈ_DEC_) of rate responses plotted as percentages of their respective control values (Wilcoxon test, p=0.563 and p=1.000, respectively).

### Effects of electric stimulation of glutamatergic afferents on the activity of the PN

We next explored the effect of activating ionotropic glutamate receptors of PM cells under conditions closer to a natural physiological context by electrically activating its innervation fibers. The strength of stimulation was adjusted to produce transient accelerations of the PN discharge (Fig. 6A, B upper trace) which were very similar in amplitude to EOD modulations triggered by the activation of the command neuron for escape behavior (Comas and Borde 2010; Curti et al. 1999, 2006). Rate responses typically peaked at the interspike interval I_1_ (see Methods), although in some cases the maximum acceleration was observed in I_0_. Overall, the mean peak amplitude of responses was 9.9±6.2 Hz and represented an increase of 59.0±29.3 % with respect to the pre-stimulus PN discharge rate. The peak of the responses was followed by a slow return to the pre-stimulus rate with a time constant (τ_DEC_) of 109.6±26.8 ms (n=14).

**Figure 6.**
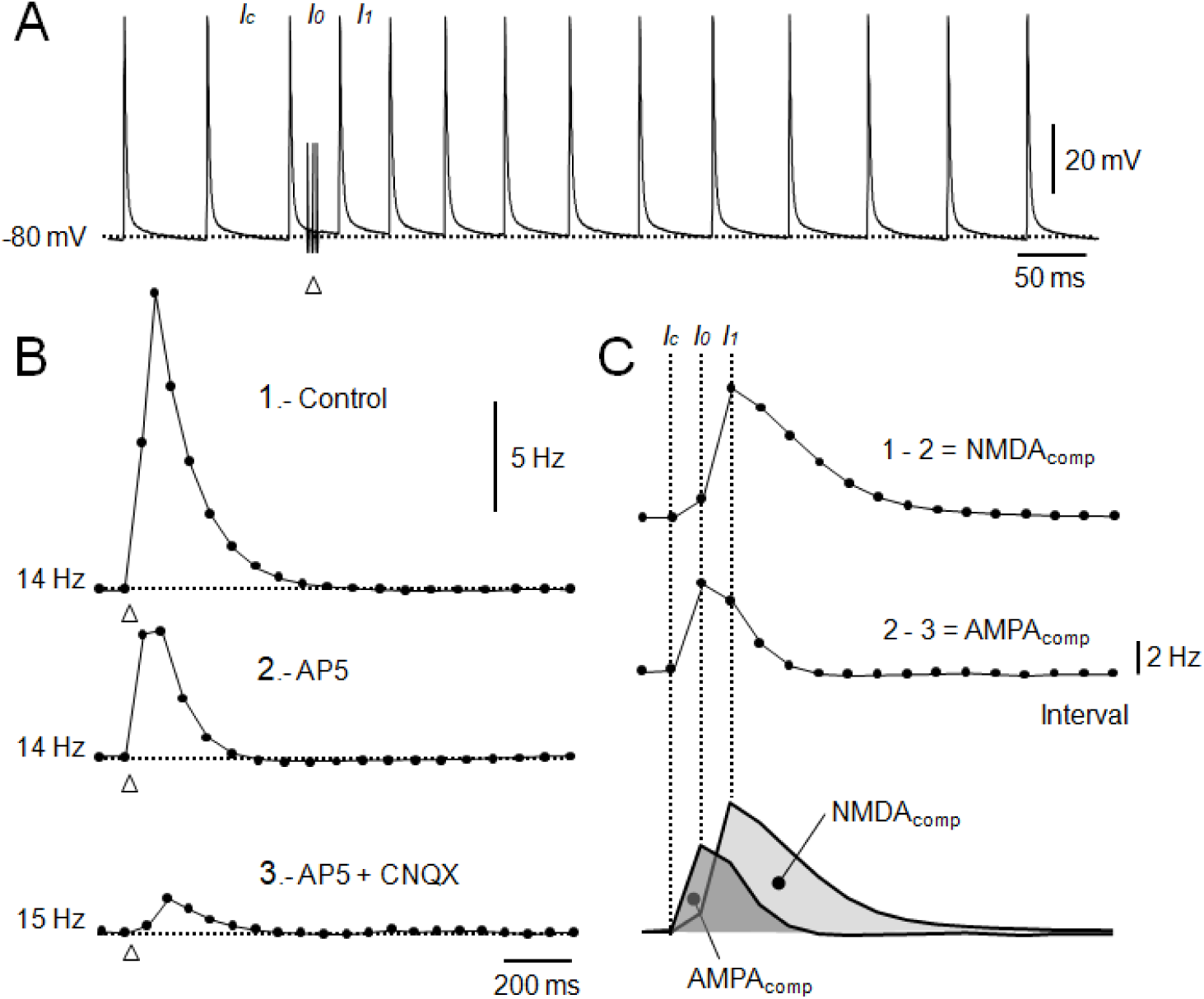
NMDA and AMPA components of the PN rate responses to synaptically released glutamate. **A.** Intracellular recording of a R-cell obtained before and after the stimulation of afferent fibers to the PN (3 stimuli @ 250Hz) in ACSF. The interval at which the stimulus is applied (I_0_) is preceded by I_C_, the control interval, and is followed by the interval at which the amplitude of rate responses usually reach its maximum (I_1_). The number at left represents the membrane potential indicated by the dotted line. **B. Top.** Plot of instantaneous frequency vs time of the recording illustrated in A (1.- Control). **Middle.** Plot of the rate response obtained after adding AP5 (50 μM) to the ACSF (2.- AP5). **Bottom.** Plot of the rate response obtained after adding CNQX (20 µM) to the AP5-containing solution (3.- AP5 + CNQX). Numbers at the left represent the pre-stimulation basal rate of discharge indicated by the dotted lines. In A and B, white arrowheads indicate the moment of afferent fibers stimulation. **C.** Plots of frequency vs. interval number of the point-by-point subtractions of the plots illustrated in B. **Top.** Component of the rate response due to activation of NMDAR (NMDA_comp_) obtained as the plot 1 minus the plot 2. **Middle.** Component of the rate response due to activation of AMPAR (AMPA_comp_) obtained as the plot 2 minus the plot 3. **Bottom.** NMDA_comp_ (light gray) and AMPA_comp_ (dark gray) superimposed in order to compare their respective time courses. Identification of intervals I_C_, I_0_ and I_1_ are indicated as vertical dotted lines as a reference.

**Figure 7.**
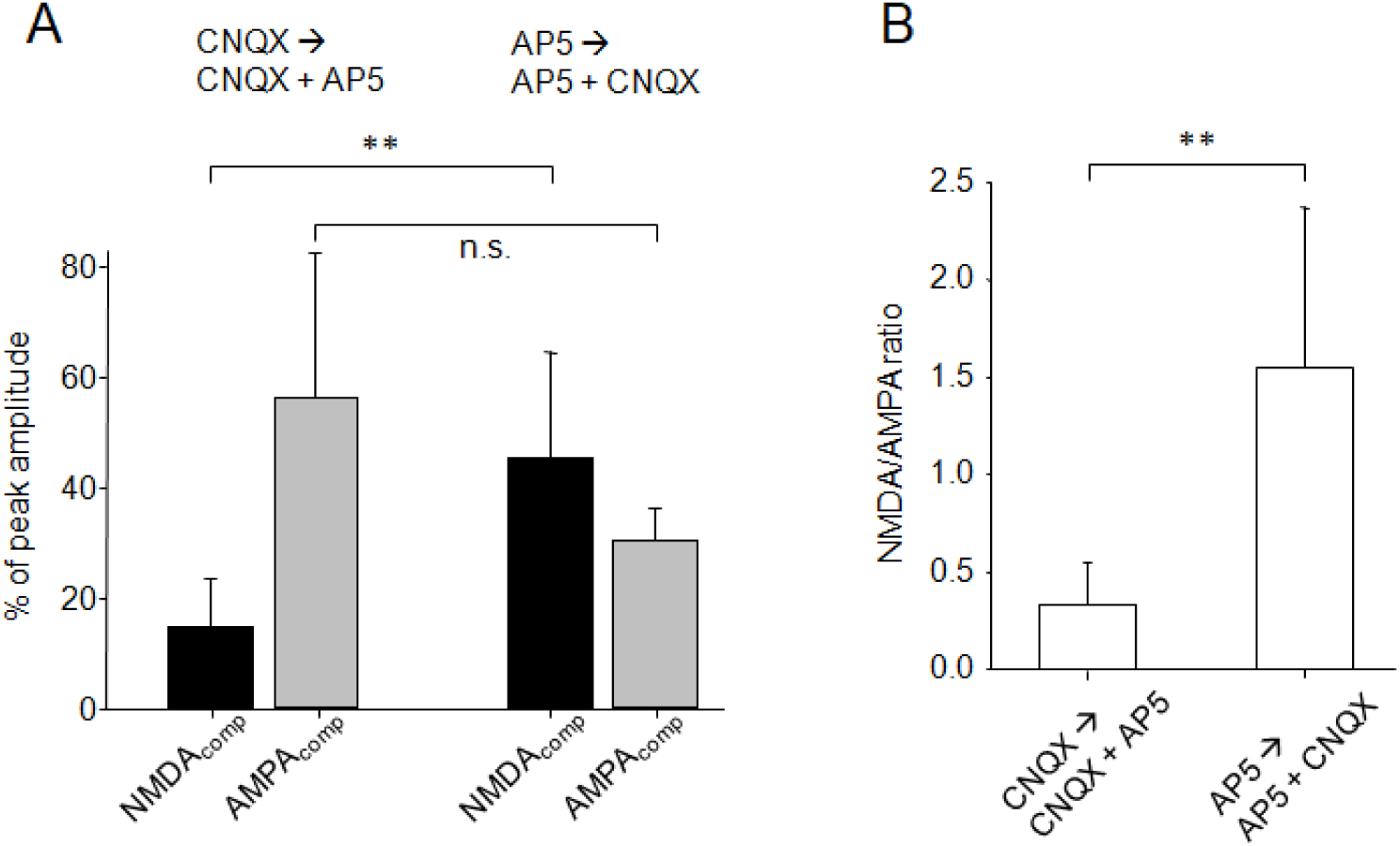
Effect of the sequence of application of blockers on the magnitude of NMDA_comp_ and AMPA_comp_ of rate responses to synaptically released glutamate. **A.** Bar plots of the mean (± SD) NMDA_comp_ (black) and AMPA_comp_ (gray) estimated by adding the blockers to the ACSF following the sequence CNQX→CNQX+AP5 (n=7) and the sequence AP5→AP5+CNQX (n=5). NMDA_comp_ (Student *t*-test, p=0.004) and AMPA_comp_ (Mann Whitney *U*-test, p=0.149). **B.** Bar plot of the mean (±SD) NMDA/AMPA ratio calculated from data represented in A for the CNQX→CNQX+AP5 sequence and the AP5→AP5+CNQX sequence (n=7 and n=5, respectively, Mann Whitney *U*-test, p=0.005).

The amplitude and the time course of accelerations of the PN discharge evoked by activation of PPn axons were specifically affected by ionotropic glutamate receptor blockers (Fig. 6B). Blocking NMDARs with AP5 (50 µM) reduced the amplitude and the duration of the response (Fig. 6B, middle trace) and the remnant acceleration was further reduced by blocking AMPARs through the addition of CNQX (20 µM) to the ACSF (Fig. 6B, lower trace). The representative example shown in Fig. 6C illustrates the procedure followed to estimate the relative contribution of PM-cell NMDARs and AMPARs to the rate response evoked by electrical stimuli. The point-by-point subtraction of the AP5 response from the control response allowed us to estimate the AP5-sensitive component of the acceleration likely due to the activation of NMDARs (NMDA_comp_, Fig. 6C, upper trace). In turn, subtraction of the response obtained after AP5 and CNQX from the AP5 response reveals the component due to the activation of AMPARs (AMPA_comp_, Fig. 6C, middle trace). Characteristically, the NMDA and AMPA components differed in their contribution to the total increase in the rate of the PN discharge evoked by activation of PPn axons. In average, activation of NMDAR contributed with a 45.5±19.1 % and AMPAR with a 30.4±5.9 % of the peak amplitude of the rate response (n=5, Fig. 7A, right bars). NMDA and AMPA components also differed in their duration and time course (Fig. 6C, lower trace). The AMPA_comp_ typically peaked during I_0_ and exhibited a faster time course than the NMDA_comp_ which reached its maximum during I_1_ and showed a larger duration. As expected by the different time course of both components, blockers affected the amplitude of the responses at I_0_ and I_1_ to a different extent. The ratio of the amplitude of the response (AR) at I_1_ and I_0_ (AR_I1_ / AR_I0_) was reduced by AP5 (from 0.77±0.77 to 0.49±0.32, Wilcoxon test, n=5, p=0.055) whereas the ratio was increased by CNQX (from 1.78±1.37 to 2.85±2.57, paired *t* test, n=8, p=0.013). Consistently, under AP5 and CNQX, a small (10-26 %) increase in the PN discharge rate in response to PPn axons stimulation was still observed (Fig. 6B, lower trace).

The above data demonstrate that NMDAR and AMPAR of PM-cells are activated by glutamate released from electrically activated innervating fibers. However, these data are not optimal for assessing whether postsynaptic effects due to NMDAR activation are dependent on AMPAR activation as it normally occurs at AMPA-NMDA transmitting glutamatergic synapses. If this was indeed the case, the magnitude of the NMDA_comp_ (AP5-sensitive) would depend on the specific order in which ionotropic glutamate receptor blockers were added to the bathing solution. We reasoned that if part of the NMDAR-mediated response was strictly dependent on the activation of AMPARs of PM-cells (co-activation of co-localized AMPAR and NMDAR), the magnitude of the NMDA_comp_ would be smaller when AP5 was added to the ACSF containing CNQX (CNQX→CNQX+AP5 sequence) compared to the one obtained adding CNQX to a solution that contains AP5 (AP5→AP5+CNQX sequence).

To test this possibility, we carried out a series of experiments to estimate the NMDA and AMPA components of rate responses in different conditions. In the first experiment AP5 was added to the slice which was being perfused with CNQX (indicated as CNQX→CNQX+AP5) and in the second experiment CNQX was added to the slice that was being perfused with AP5 (AP5→AP5+CNQX). When administration of blockers followed the former sequence, the NMDA_comp_ was indeed smaller than the one estimated with the later sequence (Fig. 7A, black bars). Overall, the mean estimated NMDA_comp_ of responses represented 14.9±8.8 % of the control response when AP5 was added to the ACSF containing CNQX and 45.5±19.1 % when the order was inverted (n=7 and n=5, respectively, Student *t* test, p=0.004). The fact that the NMDA_comp_ was consistently smaller under AMPAR blocking by CNQX suggests that part of NMDA_comp_ can be detected if and only if AMPARs of PM-cells are co-activated with NMDARs. The magnitude of estimated AMPA_comp_ was also affected by the order of blocker administration but to a lesser extent (Fig. 7A, gray bars). When blockers were applied following the CNQX→CNQX+AP5 sequence the AMPA_comp_ was 56.3±26.3% of the control response and was 30.4±5.9 % with the AP5→AP5+CNQX sequence (n=7 and n=5, respectively, Mann-Whitney *U* test, p=0.149). The overall results are summarized in Fig. 7B by comparing the NMDA/AMPA ratios using both sequences of administration of blockers. In experiments applying blockers with the CNQX→CNQX+AP5 sequence the mean NMDA/AMPA ratio was 0.33±0.22 whereas the AP5→AP5+CNQX sequence consistently result in larger NMDA/AMPA ratios with a mean value of 1.55±0.82 (n=7 and n=5, Mann Whitney *U* test, p=0.005).

### Effects of Mg^2+-^free ACSF on the NMDA/AMPA ratio

The above results indicate that at least part of the postsynaptic effects of synaptically released glutamate rely on the activation of a subpopulation of NMDARs of PM-cells that needs the simultaneous AMPAR-mediated depolarization to relieve its Mg^2+^ blockade, as in most other glutamatergic synapses (Coan and Collingridge 1985; Collingridge et al. 1988; Daw et al. 1993; Nowak et al. 1984). According to this model, we hypothesized that the dependence of NMDA/AMPA ratio on the order of administration of glutamatergic blockers should not be observed (or would be greatly reduced) if similar experiments were performed using ACSF that does not contain Mg^2+^ (nominally Mg^2+-^free). We predicted that this modified ACSF may cause a permanent relief of Mg^2+^ blockade allowing the expression of the full NMDA_comp_ of rate responses regardless of the order of blockers administration.

During preliminary experiments we noticed that the absence of Mg^2+^ (equimolarly substituted by Ca^2+^) in the medium increased the synaptic efficacy (most probably through a presynaptic mechanism; Dodge and Rahamimoff 1967) in addition to inducing changes in the magnitude of estimated NMDA and AMPA components. Consequently, to avoid a possible synaptic interfering factor, the hypothesis stated above was tested on the rate responses evoked by local application of glutamate to PM-cells instead of those provoked by electrical stimulation of glutamatergic inputs. For estimation of both components in these experiments only responses with time to peak of about 500 ms (or less) were computed in consideration of the theoretical time course of activation of different subtypes of glutamatergic receptors predicted by the diffusion model detailed in the APPENDIX.

The representative example illustrated in Fig. 8A shows the effect of the CNQX→CNQX+AP5 sequence on the rate-responses evoked by application of glutamate at PM-cells. As observed with rate responses to electrical activation of glutamatergic inputs, the magnitude of estimated NMDA and AMPA components of responses to glutamate application also depended on the order of administration of blockers (Fig. 8B). Overall, the mean estimated NMDA_comp_ of responses represented a 17.7±9.2 % of the control response when AP5 was added to the ACSF containing CNQX and 32.5±11.9 % when the order was inverted (n=5 and n=7, respectively, Student *t*-test, p=0.043) and the mean estimated AMPA_comp_ under the two conditions were 47.6±14.44 % and 31.5±15.1 % of responses, respectively (n=5 and n=7, respectively, Student *t-*test, p=0.093). The NMDA/AMPA ratio differed with the order of administration of blockers (Fig. 9A) and the difference was greatly reduced during the perfusion of slices with Mg^2+^-free ACSF (Fig. 9B). In control conditions (Fig. 9A, ACSF containing 1.2 mM Mg^2+^) the NMDA/AMPA ratio was 0.45±0.34 when the sequence of blockers was CNQX→CNQX+AP5 and was consistently larger with the inverse sequence with a mean value of 1.46±1.20 (n=5 and n=7, respectively, Student *t*-test, p=0.10). In Mg^2+^-free ACSF, irrespective of the sequence of blockers tested, there was a global increase of the NMDA/AMPA ratio suggesting a significant reduction of the voltage-dependent Mg^2+^ block of the NMDAR. Consistent with this finding, the NMDA/AMPA ratio following the CNQX→CNQX+AP5 approaches the value of the ratio with the AP5→AP5+CNQX sequence (1.80±1.05 and 2.06±1.21; n=4 and n=5, respectively, Student *t*-test, p=0.730), suggesting a lower dependence on the order of application of blockers. This was further evaluated by comparing the OD index estimated in control ACSF with the one obtained in Mg^2+^-free ACSF (Fig. 9C). Under control ACSF (1.2 mM Mg^2+^) the OD index was 2.33±0.34 and was drastically reduced to 0.15±0.68 in Mg^2+^-free ACSF.

**Figure 8.**
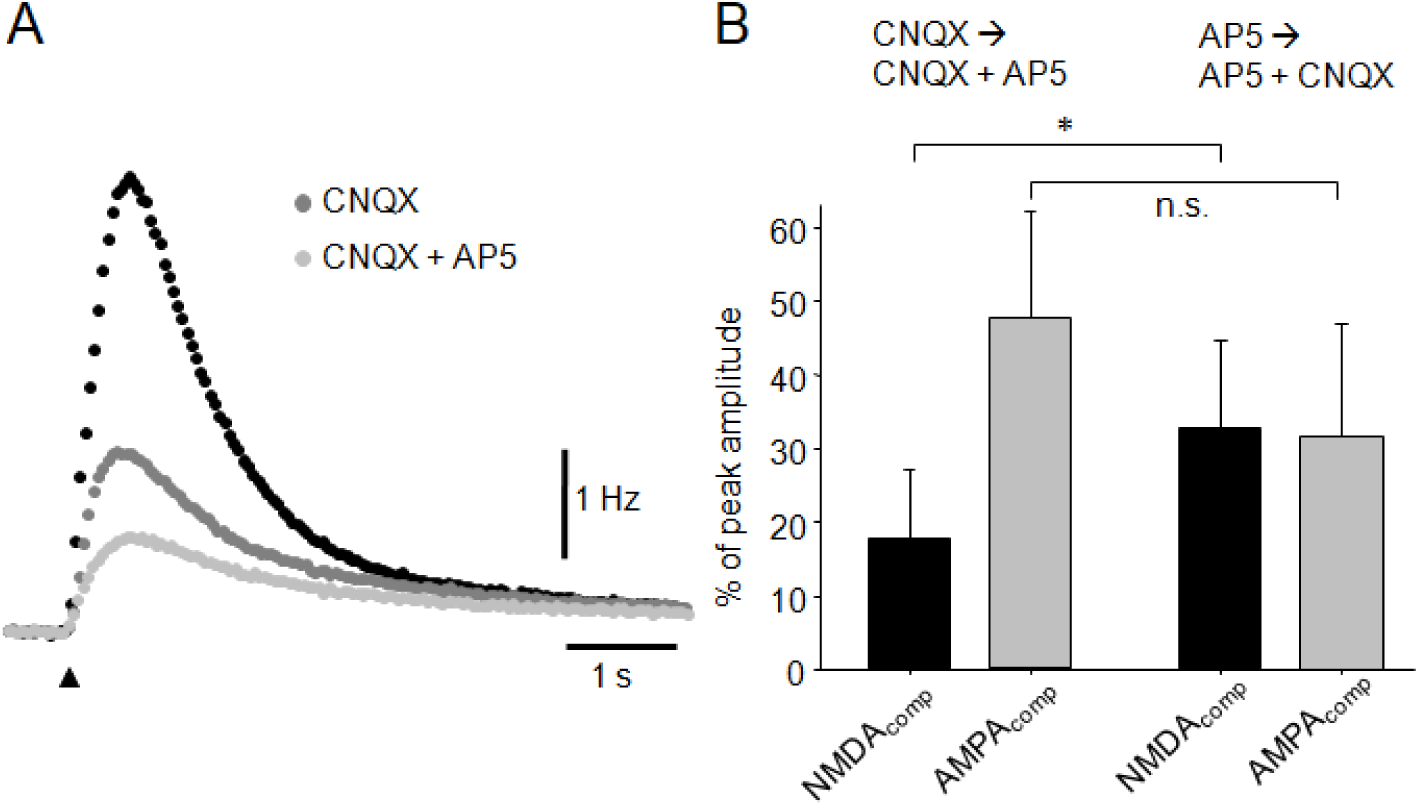
Effect of the sequence of application of blockers on the magnitude of NMDA_comp_ and AMPA_comp_ of rate responses to pressure-applied glutamate. **A.** Superimposed plots of the instantaneous frequency vs time of the rate responses to glutamate applied juxtacellularly to the PM-cells obtained before (black circles) and after adding CNQX (20 µM,) to the ACSF (CNQX, dark gray circles) and after adding AP5 (50 µM) to the CNQX-containing ACSF (CNQX + AP5, light gray circles). The black arrowhead indicates the moment of glutamate ejection. **B.** Bar plots of the mean (± SD) NMDA_comp_ (black) and AMPA_comp_ (gray) estimated by adding the blockers to the ACSF following the sequence CNQX→CNQX+AP5 (left pair of bars, n=7) and the sequence AP5→AP5+CNQX (right pair of bars, n=5). NMDA_comp_ (Student *t*-test, p=0.043) and AMPA_comp_ (Student *t*-test, p=0.093).

**Figure 9.**
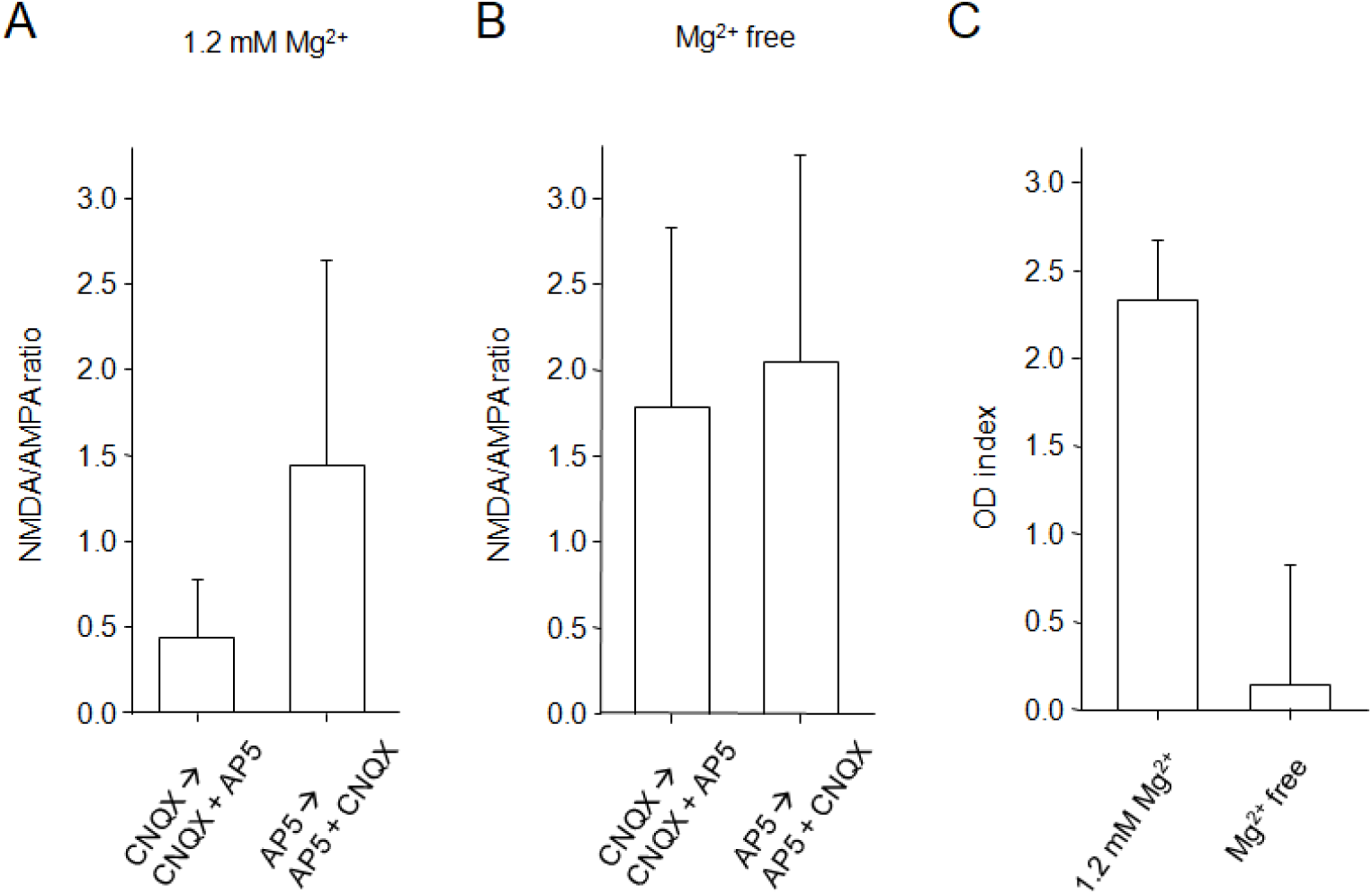
Effects of Mg^2+^ free ACSF on the blockers sequence-dependence of the NMDA/AMPA ratio of rate responses to pressure-applied glutamate. **A**. Bar plot of the mean (±SD) NMDA/AMPA ratio in ACSF (1.2 mM Mg^2+^) after adding the blockers following the CNQX→CNQX+AP5 sequence (n=7) and the AP5→AP5+CNQX sequence (n=5; Student *t*-test, p=0.10). **B.** Same as A but in a nominally free Mg^2+^ ACSF (Mg^2+^ free). CNQX→CNQX+AP5 sequence (n=4) and the AP5→AP5+CNQX sequence (n=5; Student *t*-test, p=0.730). **C**. Bar plot of the OD index (± estimated SD) under the perfusion of control (1.2 mM Mg^2+^) and nominally free Mg^2+^ ACSF (Mg^2+^ free).

## DISCUSSION

Although the mechanisms underlying the extrinsic modulations of pattern-generating networks have been well established in invertebrates, most of their synaptic and cellular basis in vertebrates are not yet completely understood (Daghfous et al. 2016; Grillner and El Manira 2020; Marder et al. 2014). The electromotor system of pulse-type gymnotiform electric fish has proven to be a well-suited vertebrate preparation to investigate these mechanisms (Borde et al. 2020; Caputi et al. 2005). In this study, we used an *in vitro* reduced preparation of the PN to do an in depth electrophysiological and pharmacological analysis of synaptic mechanisms involved in transient neurotransmitter-mediated modulations of the electromotor CPG in *G. omarorum*. Among gymnotiform fish, this species the most suitable for the study of these mechanisms since the rhythmic layer of the electromotor CPG is apparently the exclusive cellular target of the descending glutamatergic inputs. Our main findings include the following: (1) PM-cells are endowed with diverse functional glutamate receptor subtypes (NMDA, AMPA and metabotropic), while in R-cells these receptors are absent, a finding that indicates a distinctive, species-specific, distribution of glutamate receptors subtypes on PN neurons; (2) NMDAR of PM-cells are insensitive to specific blockers of receptors containing the GluN2B subunit usually associated with long lasting postsynaptic effects at only-NMDA synapses suggesting species-specificity in the mechanisms of control of PN discharge at the molecular level; and (3) synaptically released glutamate by descending inputs to the PN activates both AMPAR and NMDAR subtypes of PM-cells revealing a probable functional role of transmitting AMPA-NMDA synaptic contacts in the control of the electromotor CPG in gymnotiform fish.

### R-cells lack gutamate receptors while PM-cells neurons express different gultamate receptor subtypes

In gymnotiform fish, activation of glutamatergic inputs directed to R-cells -the critical component of the patterning layer of the electromotor CPG-trigger the emission of social electric signals that imply profound modifications of the output pattern due to suprathreshold activation of these cells through NMDAR or AMPAR (Kawasaki and Heiligenberg 1989, 1990; Quintana et al. 2011, 2014; Spiro 1997;Spiro et al. 1994). Previous work in *G. omarorum* failed to obtain any evidence suggesting there is direct suprathreshold activation of R-cells by glutamate agonists (Comas et. al 2019; Curti et al. 1999), even though these fish are capable of emitting social electric signals during agonistic encounters with distorted EOD waveform (Batista et al. 2012; Comas et al. 2019). In those studies, however, the role of subthreshold activation of R-cells -which is virtually undetectable during EOD recordings-could not be ruled out. Here, we could analyze that possibility by performing intracellular recordings of R-cells during the juxtacellular application of glutamate agonists to these cells and we show that glutamate, AMPA and NMDA were indeed ineffective in modifying either its membrane potential or its firing. This demonstrates that R-cells in *G. omarorum* lack functional glutamatergic receptors and provides additional evidence that illustrates the diversity of neural designs for the control of the electromotor CPG that have evolved in gymnotiform fish.

Juxtacellular application of glutamate agonists to PM-cells evoked transient accelerations whose amplitude and time course varied with the specific agonist applied. Our data confirm previous results obtained *in vivo* (Curti et al. 1999) indicating that, in *G. omarorum*, PM-cells are endowed with AMPAR, NMDAR and mGluR. In our experiments, accelerations produced by the pressure-applied glutamate represented an increase in the PN discharge rate of about 35 % of control the basal pre-ejection rates. According to the slight modification of the PN discharge rate caused by the injection of long lasting depolarizing current pulses (∼20 mV depolarization in Fig. 2A), a considerable depolarization during the accelerations caused by glutamate applied to the PM-cells was expected. However, rate responses to pressure applied glutamate consistently associated low amplitude depolarizations of PM-cells of at most 2.3 mV. The apparent discrepancy between the amount of depolarization and the resulting change of the PN discharge rate evoked by glutamate can be reasonably accounted for by the fact that PM-cells are electrotonically coupled (Bennett et al. 1967) and the magnitude of PN rate responses would depend not only on the net effect of the activation of glutamate receptors on the membrane potential of each PM-cell but also on the number of neurons actually activated and the timing of such activation. As proposed by Getting (1974), synchronous synaptic inputs to several electrotonically coupled postsynaptic cells (distributed input) may result in an apparent increase in the postsynaptic input resistance and time constant due to an effective reduction in the shunting of current through the junctions. This may lead to a more effective synaptic transmission producing greater postsynaptic effects in the population of coupled cells. At the PN, the more PM-cells receive synchronous depolarizing inputs the greater will be the evoked acceleration of the PN discharge (see APPENDIX). Activation of metabotropic glutamate receptors involving the modulation of a yet unknown membrane conductance involved in the pacemaker discharge but without significantly changing the MPL at PM-cells may be an additional contributing factor.

As expected by the different kinetics of ionotropic glutamate receptor subtypes (Traynelis et al. 2010), responses to AMPA were relatively faster than those to NMDA. Responses to *trans*-ACPD were even much slower than those produced by ionotropic agonists. Evidence indicating the existence of AMPAR and NMDAR in PM-cells in pulse and wave type gymnotiforms has been reported before (Caputi et al. 2005; Dye et al. 1989; Kawasaki and Heiligenberg 1990; Quintana et al. 2014). However, as confirmed in the present work, *G. omarorum* is still the sole species in which the existence of mGluR subtypes in PM-cells has been demonstrated. In this species, the motor command for the motor escape behavior -a single action potential at one identifiable reticulospinal neuron: the Mauthner cell-triggers an abrupt and prolonged increase of the EOD rate (up to 5 s) thanks to the co-activation of NMDAR and mGluR of PM-cells through a paucisynaptic neural pathway (Comas and Borde 2010; Curti et al. 1999, 2006; Falconi et al. 1997,). The long duration of this electromotor behavior triggered by a single command action potential seems to exploit the slow kinetics of glutamate receptors involved in the abrupt and long lasting activation of PM-cells. These long lasting glutamatergic postsynaptic effects mediated by the activation of NMDAR but without the co-activation of AMPAR (only-NMDA synapses) pointed to a critical role of NMDAR variants with slow kinetics and low voltage-dependence in the glutamatergic control of the PN activity. The NMDAR containing GluN2B subunits -an NMDAR variant reported as mediating only-NMDA synaptic contacts-exhibited these distinctive properties (Cull-Candy and Leszkiewicz 2004). In the present study, we show that ifenprodil, a noncompetitive NMDAR antagonist with a high selectivity for the GluN2B subunit (Williams 1993), was ineffective in modifying both the amplitude or the time course of rate responses to pressure-applied glutamate. This strongly suggests that the NMDAR containing GluN2B subunits are not involved in the glutamatergic control of the PN in this species. In *Apteronotus leptorhynchus*, a related gymnotiform species, molecular and electrophysiological data indicate that GluN2B subunits are expressed in most brain regions including the PN and that these subunit are capable of forming NMDAR whose core functional aspects are similar in fish and in mammals (Harvey-Girard and Dunn 2003; Harvey-Girard et al. 2007). However, *A. leptorhynchus* and *G. omarorum* represent two phylogenetically distant gymnotiform species (Crampton 2011; Hopkins 2009) that may exhibit species-specific differences in the neural underpinnings of electromotor displays as has been found among related gymnotiform species (Borde et al. 2020). It is therefore reasonable to expect that this diversity also involves the mechanisms of control of PN discharge at the molecular level by expressing different NMDAR subtypes with different functional characteristics.

Although the differences in the kinetics of synaptic currents mediated by AMPAR, NMDAR and mGluR (Traynelis et al. 2010) are consistent with the different time course of rate responses evoked by juxtacellular application of agonists, in our experiments those differences may also be due to other possible contributing factors. In fact, the time course of responses to the pressure-applied glutamate may be influenced not only by the relative contribution of the kinetics of AMPAR-, NMDAR- and mGluR-mediated postsynaptic currents, but also by the time course of diffusion of glutamate in the extracellular space and the differences in the EC50 values for glutamate of each receptor subtype (see APPENDIX).

### Functional role of glutamate via PM-cells ionotropic receptors

The present study reveals, for the first time in pulse-type gymnotiform fish, that synaptically released glutamate consistently activates AMPAR and NMDAR of PM-cells. This is a new finding that demonstrated that AMPARs of PM-cells, which have been described in several gymnotiform species by locally applying glutamate agonists (Curti et al. 1999; Quintana et al. 2014), can be activated by glutamate released from synaptic terminals.

It has been reported that an increase in glutamate concentration in the synaptic cleft at silent synapses (only-NMDAR) produced during certain forms of synaptic plasticity or by specific patterns of presynaptic activation, may recruit an AMPAR component converting a silent synapse into a functional transmitting one (Voronin and Cherubini 2004). However, the amplitude of the rate responses to synaptically released glutamate observed in the present study suggests that this possibility is unlikely. In fact, the amplitude of these responses are comparable to those elicited *in vivo* by Mauthner cell activation that provoke a massive recruitment of glutamatergic PPn and even so it is not possible to detect an AMPAR component of rate responses (Curti et al. 1999, 2006). This suggests that the pattern of electrical stimulation of glutamatergic axons used in our experiments -that mimicked the pattern observed during the Mauthner cell-dependent electromotor behavior (see Methods)-was indeed in the physiological range and provoked rate responses with specific AMPAR components. Moreover, preliminary data obtained *in vivo* suggest that AMPAR of PM-cells can be recruited by presumed low levels of presynaptic glutamatergic activity. We have observed that resting variability of the EOD interval significantly decreases after the application of CNQX solutions to the PN suggesting specific spontaneous activation of glutamatergic synapses involving AMPAR (unpublished results).

In this study we found that a substantial part of the total NMDAR-mediated responses (near 67 % of the total NMDA_comp_) is strictly dependent on the activation of AMPARs of PM-cells. This suggests the co-activation of the two receptor subtypes that should be co-localized at the postsynaptic membrane. The fact that the dependence of the NMDA/AMPA ratio of rate responses on the sequence of application of specific blockers (OD index∼2.33) is almost absent in nominally Mg^2+^-free ACSF (OD index∼0.15) supports the aforementioned idea. Overall, our data suggest that at least part of the postsynaptic effects of synaptically released glutamate, rely on the activation of a subpopulation of NMDAR of PM-cells that needs the simultaneous AMPAR-mediated depolarization to relieve its Mg^2+^ blockade. This is a functional trait that characterizes the most classical transmitting glutamatergic synapse (Coan and Collingridge 1985; Collingridge et al. 1988; Daw et al. 1993; Nowak et al. 1984; Traynelis et al. 2010). Despite the large amount of work aimed at characterizing the glutamatergic innervation of the PN in gymnotiform fish (Comas et al. 2019; Curti et al. 1999; Dye et al. 1989; Juranek and Metzner 1997, 1998; Kawasaki and Heiligenberg 1989, 1990; Kawasaki et al. 1988; Keller et al. 1991; Quintana et al. 2011, 2014; Spiro 1997; Spiro et al. 1994) demonstration of glutamatergic AMPA-NMDA transmitting contacts on PN neurons has remained elusive. The present study is the first report that provides evidence indicating that the control of the activity of the electromotor CPG in pulse-type gymnotiform fish includes descending PPn axons contacting PM-cells through common transmitting glutamatergic synapses (Fig. 10).

**Figure 10.**
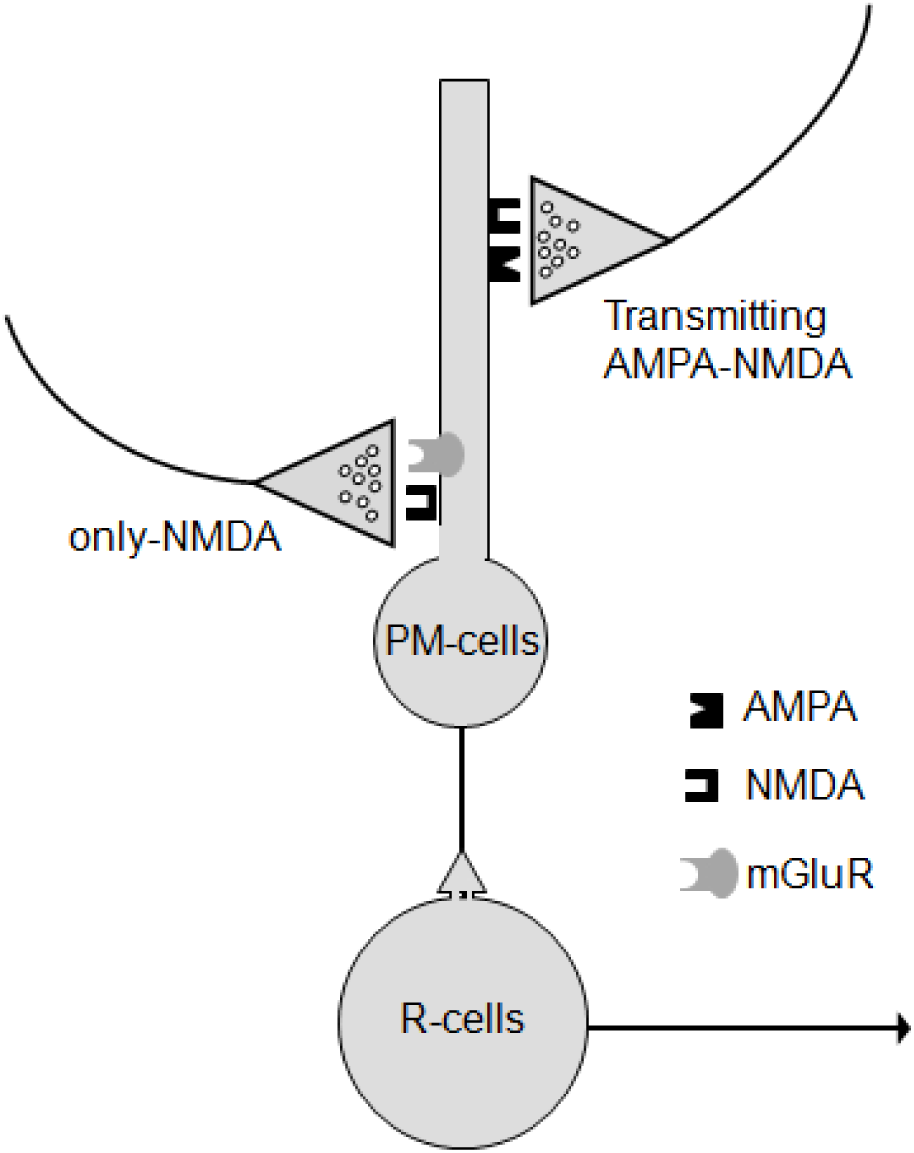
Schematic drawing of the current view of the glutamatergic control of the central component of the electromotor CPG in *Gymnotus omarorum*. Two electrotonically coupled neuronal types, PM-cells and R-cells, and different glutamate receptors subtypes are represented. The diagram summarizes our interpretation of the present data (see Discussion). The key for glutamatergic receptors are depicted in the inset.

Although the NMDA_comp_ of the rate responses to synaptically released glutamate appears to involve mainly AMPA-NMDA synaptic contacts on PM-cells there was a significant component of responses (33 % of the total NMDA_comp_) that persists under complete AMPAR blockade by CNQX. According to this, for this NMDA_comp_ to be manifest, preceding AMPA-mediated depolarizations are not necessary. This component most probably represents the activation of postsynaptic receptors at only-NMDAR synapses. But which could be the mechanism that relieves the blocking by Mg^2+^ of the NMDAR in these synapses? As discussed above, the possibility of the involvement of NMDARs with low voltage-dependence -NMDAR containing GluN2B subunits (Cull-Candy and Leszkiewicz 2004) is unlikely. Two alternative (but not mutually exclusive) mechanisms may account for the apparent low voltage-dependence of NMDAR at these synapses. First, only-NMDAR synapses in *G. omarorum* may involve receptor variants that may conduct at near resting membrane potentials as described in goldfish Mauthner cell (Wolzson et al. 1997) and in the *Apteronotus* electrosensory system (Berman et al. 1997). Second, Mg^2+^ blockade of NMDAR at these synapses may be reduced as PM-cells are spontaneously depolarizing neurons which regularly fire action potentials that may relieve (at least partially) this blockade. The latter possibility implies that only-NMDAR synaptic contacts would be located at a cellular compartment electrotonically close to the site of generation of action potentials (probably the soma). Classical AMPA-NMDA transmitting synapses, in turn, would be segregated to subcellular locations less affected by the spontaneous rhythmic membrane depolarization, for example, at the thin and long apical dendrite described by Trujillo-Cenóz et al. (1993).

According to the available evidence, PPn glutamatergic inputs to the PN neurons, which organize most of the behaviorally relevant EOD modulations in gymnotiforms, likely represent an heterogeneous population of terminals (see for a review Caputi et al. 2005). This population comprises: i, only-NMDAR synapses on both PM- and R-cells mediating accelerations and sudden interruptions of EODs, respectively, and ii, only-AMPAR synaptic contacts on R-cells whose activation is responsible for the emission of chirps (Juranek and Metzner 1997, 1998; Kawasaki and Heiligenberg 1989, 1990; Kawasaki et al. 1988; Keller et al. 1991; Quintana et al. 2014; Spiro 1997). Only-AMPAR synapses have also been characterized in mammals (Asztely et al. 1997). In *G. omarorum*, in turn, the glutamatergic control of PN discharges most probably also includes a heterogeneous population of terminals exclusively contacting the PM-cells (Fig. 10). This population includes only-NMDAR and AMPA-NMDA synapses although the existence of only-AMPAR synapses could not be ruled out. While activation of only-NMDAR synapses underlies the electromotor behavior triggered by the activation of the Mauthner cell (Curti et al. 1999, 2006), the specific EOD modulation mediated by activation of transmitting AMPA-NMDA synaptic contacts has not yet been determined. It is generally accepted that increases in EOD rate result in an enhancement of the electrosensory sampling fish perform of the environment to cope with fast changing surrounding conditions (Caputi et al. 2003; Comas and Borde 2010; Hofmann et al. 2013). EOD rate can also be modulated to avoid interfering detrimental effect of EODs generated by a nearby conspecific (Jamming Avoidance Response, JAR; Caputi et al. 2005; Heiligenberg 1991). In pulse-type fish, the interference is avoided by abruptly accelerating the EODs just before the occurrence of a coincident discharge and lasting a few tens of milliseconds (Capurro et al. 1998, 1999). Long lasting increases (seconds) in electrosensory sampling triggered by novel stimuli or during motor activity, most probably rely on activation of only-NMDA synapses on PM-cells. AMPA-NMDA synapses, on the other hand, may be involved in the rapid adjustment of the timing of the EOD during the JAR, making the most of the faster kinetic of the AMPAR (Metzner 1993).

In conclusion, our data shed light on the synaptic mechanisms underlying the fast neurotransmitter control of a vertebrate CPG. We found that, in *G. omarorum*, the rhythmic layer of the electromotor CPG is the exclusive target for the glutamatergic descending inputs contacting the PM-cells. In this species, the modulation of the CPG mediated by fast neurotransmitters exclusively comprises transient modifications of the EOD rate. The absence of functional glutamatergic receptors at the medullary neuronal components of the pattern formation layer support the hipothesis that the emission of transient communication signals that imply changes in the EOD waveform, most probably involves the extrinsic neuromodulation of these neurons (Comas et al., 2019). Our data also indicate that the glutamatergic control of the CPG likely involves both AMPA-NMDA and only-NMDA synapses contacting pacemaker cells at different cellular compartments. Contrasting with findings in mammals, only-NMDA synapses at this CPG are also transmitting synapses and likely involve NMDAR that does not contain GluN2B subunits. Active electroreception requires a stereotyped and regular EOD, the behavioral output of the CPG, for use as an energy carrier for perception. According to the complement of postsynaptic receptors of synapses contacting PM-cells, glutamatergic descending inputs to the CPG are most probably involved in modulations displayed to preserve (AMPA-NMDA synapses) or to enhance (only-NMDA synapses) the electrolocative ability of the fish.

## APPENDIX

An accurate knowledge of the time course and the spread in the extracellular space of glutamate (Glu) ejected by pressure greatly facilitates the interpretation of the effects on the discharge rate of the PN obtained by local application of this transmitter as used in some of the above experiments (Figs. 3, 4, 5 and 8).

The spread of Glu in the extracellular space can be estimated by solving the diffusion equation when the boundary conditions are appropriate (Stone 1985). Since the distance between the tip of the electrode and the membrane of the PM-cells is significantly smaller than the distance between the tip of the electrode and the surface boundaries of the recording chamber, the diffusion process is equivalent to the release of glutamate from a source point into an infinite space. The solution of the diffusion equation with the boundary conditions described above gives the concentration of Glu (C) at any time (t) and the relative position of the electrode tip (r).

Considering the micropipette tip an instantaneous source point of Glu, since the compound is ejected at a relatively short pressure pulse (duration of 50-100 ms), located in an anisotropic medium, the concentration (C) achieved at a certain time (t) is given by:

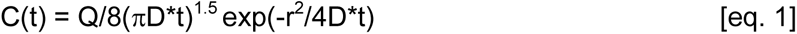

where Q (µM) is the total amount of glutamate ejected from the pipette, D* is the corrected diffusion coefficient of Glu for a tortuosity (λ) of 1.6 and an available fraction of brain tissue (α) of 0.3 obtained by applying the formula D*= (D/λ^2^) α (Nicholson et al., 1979) and r is the distance at which the concentration is calculated (with the diffusion coefficient [D] of Glu being 5.10^-6^ cm^2^. s^-1^ from Krnjevic and Phillis 1963).

An example of the time course of Glu concentration after pressure ejection at three different distances from the tip of the ejection pipette is illustrated in Fig. A1.

**Figure A1.**
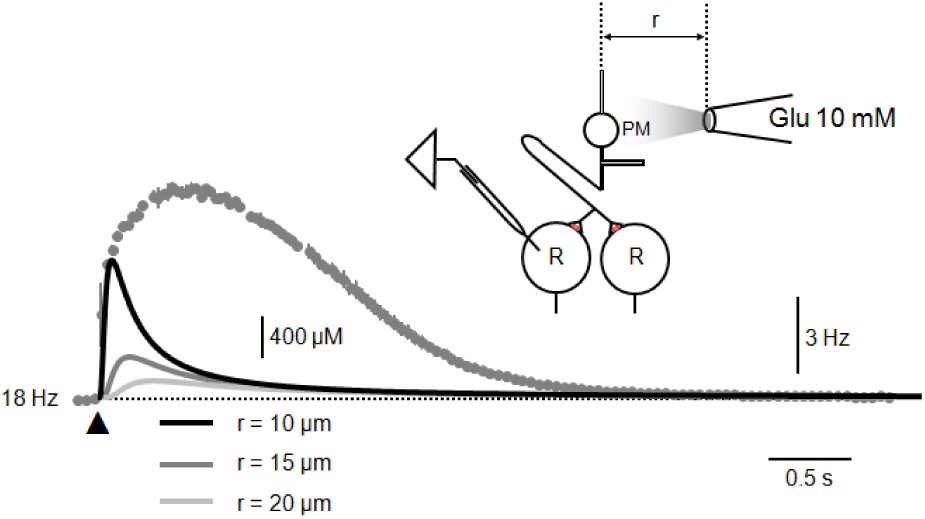
Theoretical calculations of Glu diffusion after juxtacellular pressure application and a representative response of PN discharge rate to Glu obtained during experiments. The inset in the upper right section illustrates a scheme of experimental conditions (PM, pacemaker cell; R, relay cell) that shows a recording micropipette inserted in a relay cell to monitor PN activity during Glu local applications. The superimposed plots indicate the PN discharge rate (grey filled circles) and calculated Glu concentration (lines) at three different distances from the micropipette tip vs time. Each point of the discharge rate vs time plot represents the mean (± s.d.) of 10 average responses to juxtacellular applications of Glu (arrowhead). The dotted line indicates the basal pre-ejection PN discharge rate (number on the left).

For calculations illustrated above, the ejected volume of Glu (10 mM) corresponds to the volume of a droplet of 50 µm diameter which is within the range of volumes used in our experiments.

The disparity of the time course of PN rate modulations and the calculated extracellular concentration of Glu at PM-cells membrane suggest that several other factors suggests that other factors should be taken into account to explain the time course of the responses. Since PM cells are most likely electrotonically coupled (Bennett 1971; Bennett et al. 1967; Curti et al. 2006), it is reasonable to assume that the total effect of Glu on the PN discharge rate depends on two main factors (Getting 1974):

i.- the magnitude of activation at each PM-cell that in turn relies on the concentration of Glu at the pericellular space and on the subtype and density of Glu receptors on the neuronal membrane, and

ii.- the number of neurons activated by Glu and acting on different Glu receptors and on the timing of this activation.

Assuming that the densities of NMDAR, AMPAR and metabotropic glutamate receptors (mGluR) are almost the same at PM cells and even ignoring differences in receptor kinetics, it is possible to estimate the time course of the number of cells activated by a given Glu receptors subtype according to their respective EC50 for Glu. This estimation implies the calculation of the spread of glutamate in the extracellular space by defining a theoretical sphere whose surface represents a concentration of Glu that equals the EC50 for a given receptor subtype. For the purpose of calculations, the EC50 for AMPAR (100 µM), NMDAR (3 µM) and mGluR (10 µM) were taken from Traynelis et al. (2010).

By rearranging equation 1, the time course of the radius of such a sphere can be calculated:

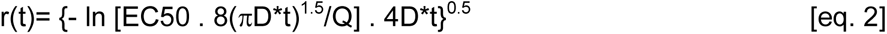

If the concentration of Glu at the surface of the sphere equals the EC50 for a specific Glu receptor subtype, it can be expected that the sphere contains neurons whose Glu receptors will be activated in a proportion higher than 50% of the available population of this receptor subtype. Neurons located out of this limit could also be activated but under a weak Glu stimulation. Therefore, following the above considerations, the numbers of cells recruited by Glu according to eq. 2 may represent an underestimation of the total number of cells actually activated.

An example of these calculations is illustrated in Fig. A2.

**Figure A2.**
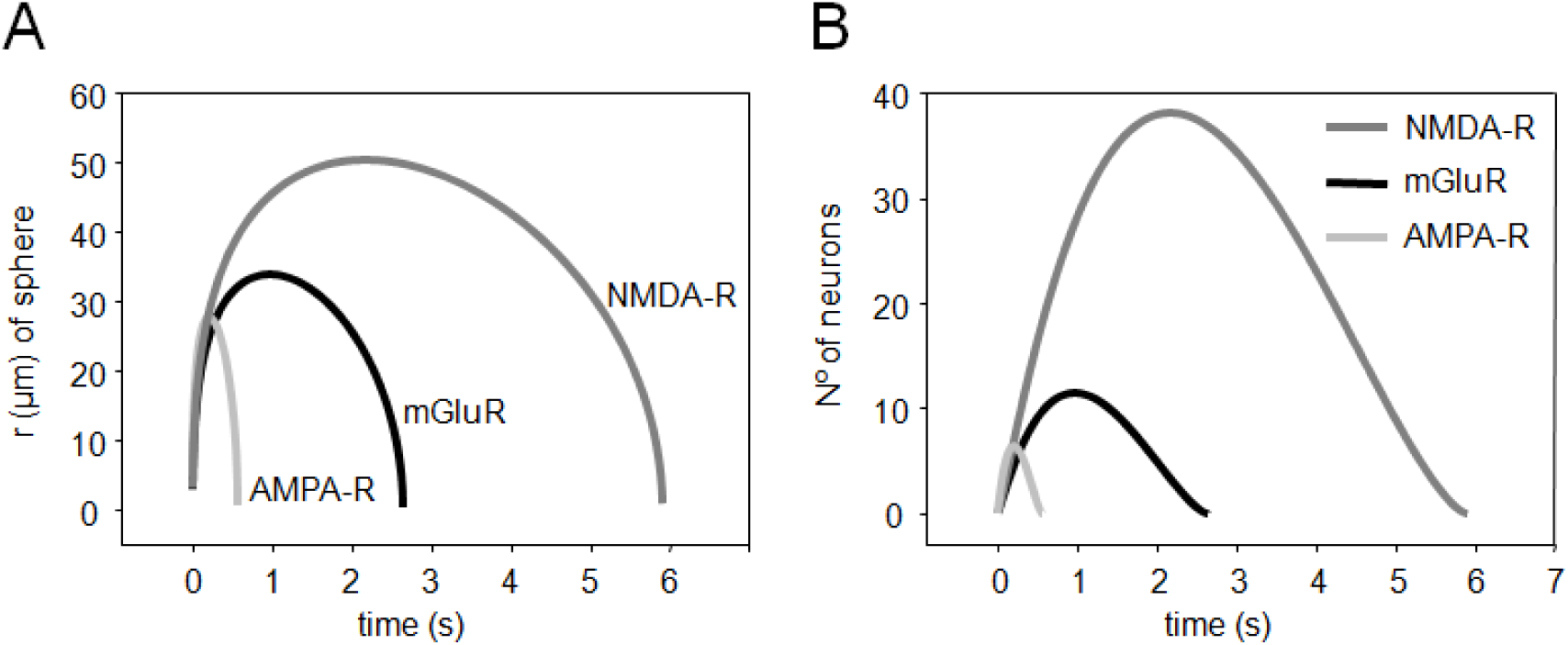
Theoretical estimation of the activation of glutamatergic receptors by the spread of glutamate in the extracellular space. A.- Plots of the radius of spheres limited by an extracellular concentration of Glu that equals the EC50 for NMDAR (dark grey), AMPAR (light grey) and mGluR (black) as a function of time. Note that the volume of tissue at which Glu extracellular concentration is able to activate different Glu receptor subtypes varies according to its relative affinity for the neurotransmitter. B.- Plot of the approximate number of cells contained within the limits of spheres illustrated in A. For this calculation, each PM-cell was assumed to be spherical (without dendrites or axons) with a diameter of about 25 µm (average diameter of real PM-cells) and to be in contact with neighboring PM-cells.

According to our calculations, a microvolume of Glu (10 mM) applied among the population of PM-cells will differently activate an heterogeneous population of PM cells which varies with time (see Fig. A2B). Immediately after Glu ejection, during the first 500 ms, a small group of cells (a maximum of 6) will be activated through a combination of AMPAR, NMDAR and mGluR. In contrast, 1 s after ejection (t=1) a significant number of PM-cells will be activated only through NMDAR and mGluR and, surprisingly, at t=2.5 an even more numerous group of PM-cells will be recruited, almost exclusively, via activation of NMDAR subtypes. This spatial sequence of PM- cells recruitment through different combinations of Glu receptor subtypes may explain, at least partially, the disparity of the time course of the effect of Glu on the rate of PN discharge and the extracellular concentration of Glu illustrated in Fig. A1.

## Acknowledgements

The authors want to thank Ana Silva and Laura Quintana for their generous comments on a preliminary version of this article. This research was supported by PDT (S/C/IF/54/090), CSIC-CAP, UdelaR and PEDECIBA

## Author contribution

V.C. and M.B. conception and design of research; V.C. performed experiments; V.C. and M.B. analyzed data; V.C. and M.B. interpreted results of experiments; V.C. and M.B. prepared figures and wrote the paper; V.C. and M.B. edited and revised manuscript and approved its final version.

## Notes

### Competing Interest Statement

The authors have declared no competing interest.

